# Mapping Alzheimer’s neuropathology signatures to the whole brain transcriptome using machine learning data-fusion

**DOI:** 10.64898/2026.08.19.745861

**Authors:** Anwesha Bhattacharya, Chloe Savignac, Liam Hodgson, Jack Stanley, Guy Wolf, Smita Krishnaswamy, David A. Bennett, Elizabeth B Binder, Danilo Bzdok

## Abstract

In Alzheimer’s disease (AD), misfolded proteins emerge across the entire brain in structured, yet not rigid, spatiotemporal patterns. Yet, a systematic bias of single-cell genomics toward sampling mostly cortical tissue limits our understanding of the whole-brain transcriptomic vulnerability to AD. Here, we develop a machine learning method to extrapolate local AD neuropathology signatures to the whole brain. By analyzing gene expression profiles of over two million cortical cells from 427 humans spanning the AD-pathology spectrum, we derive transcriptomic estimators of AD neuropathology. After extensive validations on datasets with known ground truth, we apply this framework to three million cells from 108 brain regions in the *Siletti* whole human brain atlas and derive an anticipated brain map of transcriptomic signatures indexing AD neuropathology. This interrogation of regions spanning the cortical, subcortical, and brainstem structures uncovers transcriptomic signatures associated with hyperphosphorylated tau in the medulla oblongata, dorsal raphe nucleus, and the tuberal and mammillary regions of the hypothalamus. At the cellular level, assessments of these signatures across 31 cell populations identify VGLUT1/2 expressing neurons, astrocytes, and microglia as key neuropathology-resembling populations. Within the hippocampus, pathology signatures surface in the rostral cornu ammonis (CA) subfields, particularly in the CA1 pyramidal neurons and dentate granule cells. β-amyloid-like signatures localize to the neocortex with laminar selectivity — most prominently in upper layer somatostatin+ intratelencephalic neurons (L2-L3), but also in deep layer intratelencephalic and corticothalamic neurons (L5-L6). Neocortical astrocytes and microglia exhibiting disease associated signatures similarly demonstrate a unique laminar preference. Together, this study provides the first whole human brain map of AD pathology-associated transcriptomic signals, and exposes cell type, region, and cortex layer specific vulnerabilities.

## Introduction

Single-cell genomics technologies have transformed the study of neurodegenerative disease by resolving cell type-specific molecular alterations within complex brain tissue. These approaches have revealed selective cellular vulnerabilities, disease-associated states, and regulatory programs that underlie pathogenic progression ^1^. However, most of these efforts remain constrained by tissue availability, cost, and technical complexity ^2^. As a result, studies typically focus on one brain region at a time — most often neocortical tissue ^3,4^ — and, at best, include a small a priori-selected set of cortical, often prefrontal, and medial temporal lobe regions ^5–8^. Consequently, large portions of the adult human brain remain under-explored at the cellular and molecular resolution, creating a systematic blind spot in our understanding of disease biology.

This neuroscience community’s cortex-centric sampling bias is particularly consequential for Alzheimer’s disease (AD) in which pathological lesions affect almost the entire brain in a highly selective manner. Based on clinical frameworks anchoring symptomatic AD to cognitive impairment, cortical systems that support memory (primarily hippocampus) and higher order functions (prefrontal cortex and middle temporal gyrus) have been prioritized. Neuropathological evidence, however, tells a different story. Seminal staging studies in postmortem autopsy samples have shown that subcortical and deep brain structures often exhibit pretangle lesions years before the emergence of overt cortical tangles or plaque deposits ^9–11^.

Yet the precise anatomical spread, cellular substrates, and molecular basis of the earliest events in AD remain poorly understood. In particular, small brainstem nuclei and deep subcortical structures have historically been underrepresented across many modalities of AD research, including single-cell genomics, and in vivo structural and functional imaging studies ^12–14^. Resolving initial disease-relevant vulnerabilities is critical, because therapeutic interventions are likely to be most effective before widespread pathology and clinical symptoms are manifested ^15^.

Despite neuropathological evidence supporting the need to study AD in a brain-wide setting, the lack of deep-brain, non-cortical transcriptomic datasets creates a fundamental gap in today’s data real estate. While disease-focused single-cell datasets provide rich AD-relevant phenotypes and neuropathological annotations, they are generally anatomically restricted ^16^. Conversely, the largest, whole-brain transcriptomic atlas in humans recently emerged to provide a broad cellular and spatial coverage, but it lacks matched AD neuropathology measurements. At such scales of millions of cells, generating a dedicated whole-brain AD resource would require substantial coordinated efforts and is unlikely to be feasible in the years to come.

To begin closing this gap, we developed a computational data-fusion framework that transfers disease-relevant information from AD cohorts—with matched transcriptomic, clinical and neuropathological annotations—onto the whole adult human brain. The *Siletti et al.* ^17^ snRNA-seq atlas provides the necessary transcriptomic scaffold for this mapping exercise. By data fusion of multiple complementary views of brain biology from Religious Orders Study and Rush Memory and Aging Project (ROSMAP) ^18^, SEA-AD ^19^, BRAIN Initiative Cell Census Network, and ENCODE ^20^, we augment this general atlas with transcriptome-inferred signatures of tau tangles, β-amyloid plaques, and other neuropathology markers.

By introducing novel data-fusion tools, we quantify AD-critical gene expression signatures of tissue pathology in single cells of the *Siletti* atlas sourced from 108 brain regions across cortex, subcortex, and the brainstem. Further, across 31 unique cell populations, we isolate the genes and transcriptional regulators driving the estimated regional vulnerability to pathology — providing putative therapeutic targets. Further, within the neocortex, we resolve the laminar organization of the estimated pathology signatures. Collectively, our unprecedented data science protocol enables a whole-brain, single-cell resolved map of neuropathology-associated molecular states, including regions and cell populations not yet accessible to AD single-cell profiling.

## Results

### Modelling neuropathology at single-cell resolution

We set out to characterize AD/AD and related dementia (ADRD) pathology-associated gene expression across the cellular and spatial landscape of the whole human brain. To this end, we developed a data-driven computational framework that uses interpretable machine-learning models to infer cell-level neuropathology signatures from single-nucleus RNA sequencing (snRNA-seq) data. Our pattern-learning models were trained on >2 million transcriptomes derived from the prefrontal cortex (PFC) of 427 postmortem human donors with gold standard neuropathology assessments ^21^ (dataset referred to as ROSMAP-PFC). We then used these models (pathology-estimator) to quantify neuropathology-associated signatures (pathology-proxy) in held-out donor cell transcriptomes; and even in cells from other datasets, in an across-dataset transfer-learning effort (Fig. 1; Supplementary table 1; Methods).

**Figure 1.**
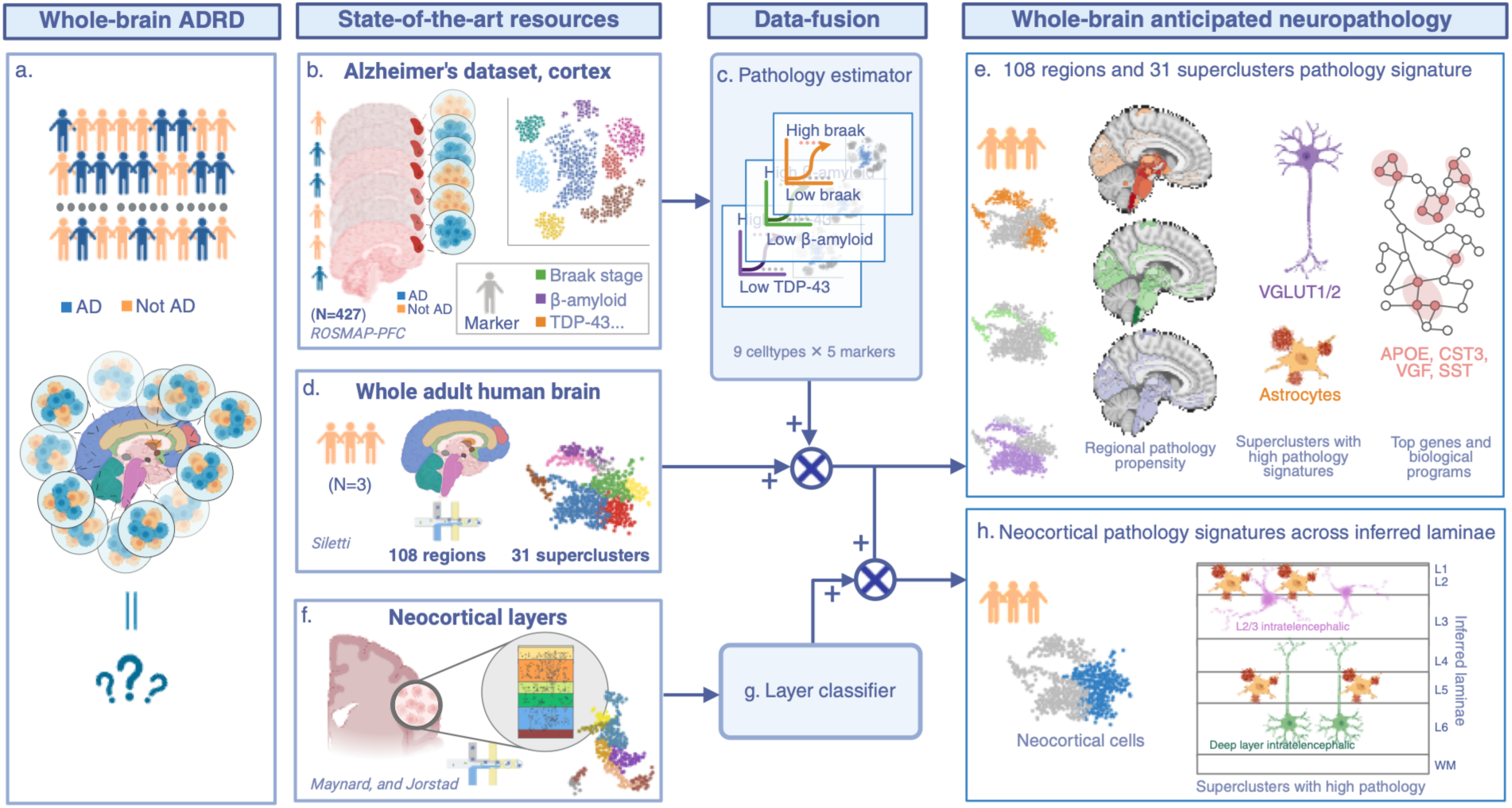
Study design. Schematic detailing the modelling framework for deriving an anticipated whole brain map of neuropathology in humans. a. Overview of the absence of a whole brain single-cell transcriptomics atlas for AD/ADRD neuropathologies. b-c. ROSMAP data from the prefrontal cortex (PFC) is used to train 45 models (9 cell types × 5 markers) to infer neuropathology signatures at the single cell level. d-e. The models are applied to a whole human brain atlas to track pathology signatures in 108 brain regions and 31 superclusters. f-g. Datasets with neocortical laminae ground truth are used to train machine learning models for layer classification in single cells. h. The layer classifiers are used to infer neocortical layer identity of cells from the whole brain atlas. Figure created using BioRender.

Our neuropathology estimators were trained on nine available cell types in ROSMAP-PFC: astrocytes (Ast), immune cells (Imm), oligodendrocytes (Oli), oligodendrocyte precursor cells (Opc), inhibitory neurons (Inh), three excitatory neuron groups (Ex1/2/3), and vascular cells (Vasc). We constructed pathology estimators for five neuropathology markers: 1) Braak stage, 2) β-amyloid (Abeta), 3) CERAD, 4) TDP-43, and 5) Lewy body. Our evaluations of these 5 pathology estimators trained independently for 9 cell types (45 models total) showed cell type-specific findings.

Across 5 neuropathology markers in 9 cell types, the trained models discriminated accurately between high and low pathology states (binarized reference pathology; Methods). Evaluations of these models on held out donors showed above-chance receiver operator characteristic area under the curve (ROC-AUC) values (Supplementary table 2-4; Methods). The maximum ROC-AUC reached up to 0.87 for telling apart high vs. low Braak (Braak stage) in Ast (average across donor-stratified cross validation folds; Methods). Further, training and evaluation under male-only, female-only, and full-cohort (i.e., including male and female donors) settings revealed a consistent dependence of predictive performance on cohort sex composition. This suggested that sex-associated transcriptomic heterogeneity was an important source of variation in our analyses. Given our target whole brain dataset (*Siletti*) comprised exclusively male donor transcriptomes (see below), we carried out all subsequent across-dataset analyses in males.

We next examined the driving genes (significantly pathology-linked genes; Methods) behind the 45 pathology estimators in further detail. Pairwise comparisons between different markers revealed low overlap, across cell types (gene effect sizes; Supplementary Fig. 1). This indicated relatively unique signatures captured by each examined cell population, for a neuropathology marker at hand.

Further probing of the transcriptomic correlates within individual cell types revealed that, Ast models were consistently the most similar across different pathologies (Kendall’s tau-b (*τ_b_*) up to 0.5 between Braak and Abeta). This prompted us to investigate the genes implicated by the 5 Ast-specific estimators. An enrichment analysis of the genes with high coefficient values from these models turned out to reflect biological processes related to protein refolding and autophagy (Methods; Supplementary table 5). All other cell types demonstrated low associations between pathologies (max *τ_b_*= 0.3 in Ex1 neurons between Braak and CERAD). Together, these observations were informative in the interpretation of subsequent analysis—which cell type- and pathology-estimators had similar transcriptomic information versus which of them captured unique axes of biological variations.

Further, we contrasted the pathology estimator models with independent control models fitted to predict age of donor (9 cell type-specific estimators; Supplementary Fig. 2c, d). Analysis of the gene-level coefficients showed low associations between the age- and pathology-estimators with the maximum average *τ_b_* being 0.06, across cell types (TDP-43 and age; Supplementary Fig. 1). This indicated that our pathology specific estimators captured molecular signatures that were distinct from general aging.

Taken together, here we showed that our cell type-specific pathology estimator models captured meaningful transcriptomic information to distinguish cells from high vs low pathology donors. We next evaluated these models on external snRNA-seq datasets to assess their generalizability to unseen donors and cells. Note that since Braak and Abeta are two primary AD neuropathology markers, we focus the reported results on these two markers in the following sections. Results from other markers are provided as Supplementary information.

### Neocortex-to-neocortex validation of neuropathology estimators

After deriving transcriptomic estimators for five neuropathological markers in the PFC, we next tested whether these models can generalize to a new cortical area. Specifically, we applied the PFC trained models from ROSMAP to an independent snRNAseq dataset from the middle temporal gyrus (MTG) of the SEA-AD cohort (>500,000 cells from 36 donors; Supplementary table 1-2; Methods).

To enable cell type-matched pathology estimation, we first assigned each SEA-AD cell to its closest ROSMAP-PFC cell type using a cell type inference model (Figure 2a; Methods). The inferred cell labels approached near complete agreement with the ground truth cell type annotations provided by SEA-AD: 18 of the 23 SEA-AD defined cell classes mapped unambiguously to a single ROSMAP-PFC cell type (e.g., all SEA-AD astrocytes mapped to ROSMAP-PFC Ast). The five excitatory neuron classes from SEA-AD were allocated to Ex1, Ex2 and Ex3. This successful alignment justified our subsequent application of the matched cell type-specific pathology estimators to each SEA-AD cell.

**Figure 2.**
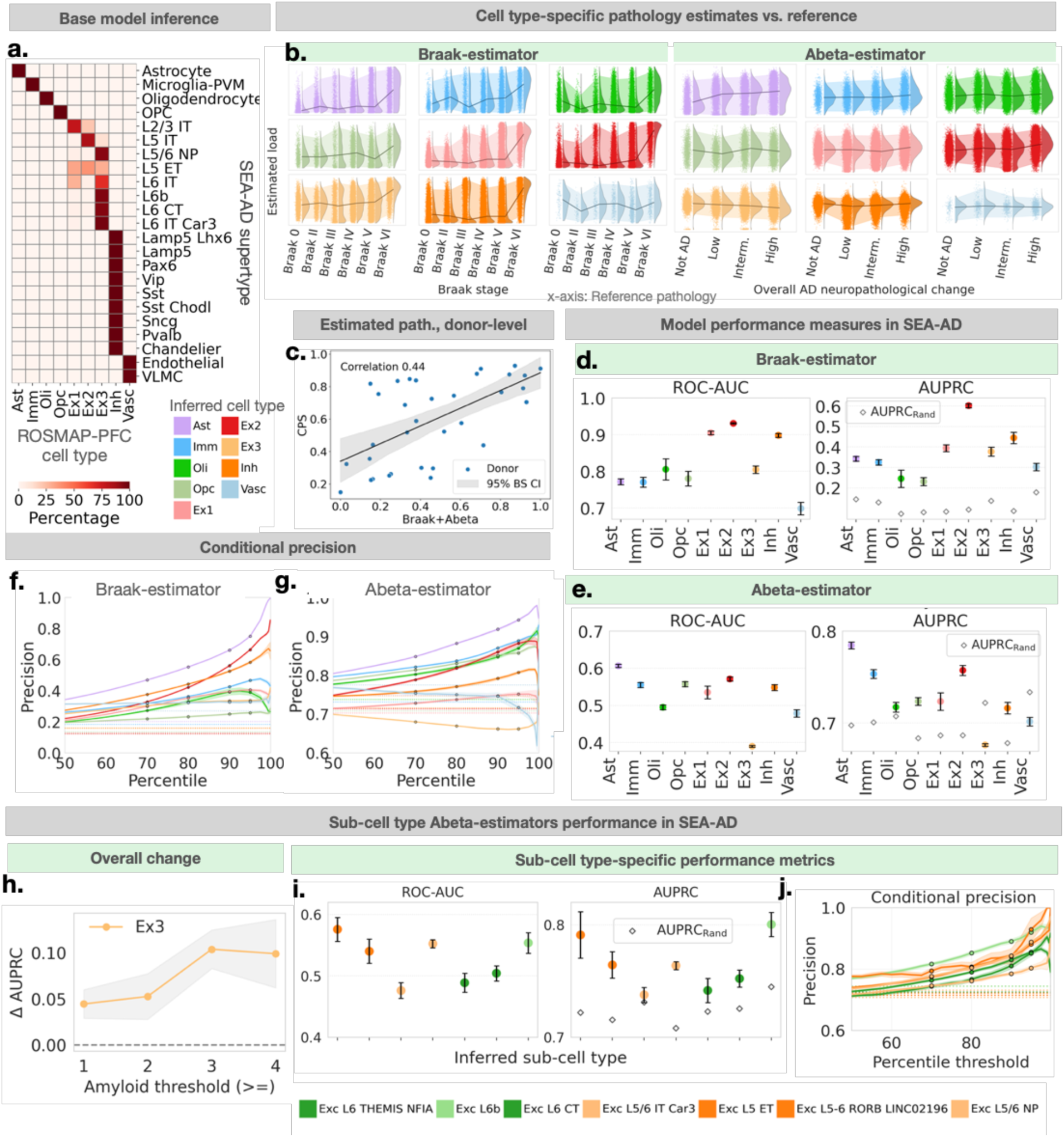
AD neuropathology estimation models transfer successfully across independent datasets. Cell-type specific neuropathology estimation models trained on ROSMAP-PFC were applied to SEA-AD single cells. a. Cell type label transfer from ROSMAP-PFC to SEA-AD cells. Each grid shows the percentage of cells from each SEA-AD supertype (x-axis) inferred to be a certain ROSMAP-PFC cell type (y-axis). b. Pathology score distributions stratified by SEA-AD reference pathology stage. Each subplot is an inferred cell type; dots represent single cells. Solid line connects median of the score distributions. Shaded regions denote 10th-90th score percentiles. Left, Braak stage (Braak); right, β-amyloid (Abeta). c. Donor-level association between estimated pathology and reference pathology (continuous pseudo-progression score; CPS). Association computed using Kendall’s tau-b. Dots represent donors; values are per-donor average across cells (normalized to [0,1]). Solid line indicates linear regression fit; shaded region denotes 5-95% confidence interval from bootstrapped models (resampled donors; n=1000 iterations). d-e. Pathology estimator model performance in SEA-AD. ROC-AUC (left) and AUPRC (right) for d, Braak, and e, Abeta. Dots denote mean, error bars indicate standard deviation (10 model refits in ROSMAP-PFC using random 50% cell subsampling). Diamonds in AUPRC panel represent baseline precision (AUPRC_Rand_=number of true positives/total number of cells). f-g. Conditional precision curves derived from estimated pathology score percentiles in SEA-AD. Solid curves connect mean precisions for each inferred cell type (10-folds; see above); shaded region denotes standard deviation. Dotted lines show baseline precision. Precision increases over the baseline at higher thresholds, except for Ex3^Abeta^ model. h-j. Ex3 neuron pathology estimators perform better with subtype-level modeling. Abeta estimators were trained for 9 ROSMAP-PFC-defined Ex3 subtypes and applied to SEA-AD Ex3 cells with matched inferred subtypes. h. ΔAUPRC=AUPRC_Ex3_subtype-model_ – AUPRC_Ex3_model_. i,j. Plots show ROC-AUC (left), AUPRC (center) and threshold-derived precision curves (right) for SEA-AD cells. Each dot (or line) represents ROSMAP-PFC cell subtype.

We assessed model performance in unseen cells from the MTG by comparing pathology-proxy scores against SEA-AD reference pathology labels (Figure 2b). Most cell types showed a monotonic increase in estimated scores with increasing measured pathology severity—showing that the model-derived pathology estimates really confirm known severity based on tissue examination in post-mortem brains. This confirmatory trend was particularly pronounced in the Braak models, which showed sharp score increases at late reference pathology stages. A donor-level assessment of aggregated Braak and Abeta estimates (averaged across all cells per donor) revealed a robust positive association with the reference continuous pseudo-progression score for AD (*τ_b_*=0.44; Figure 2c) ^19^. This indicated that donors with more advanced reference pathology systematically carried higher pathology-proxy loads.

We next evaluated model discriminative accuracies, for high vs low pathology, as measured by ROC-AUC (Figure 2e, g; Supplementary table 6; Methods). For Braak estimation, all inferred cell types achieved performance above chance level (i.e., 0.500) with Ex2^Braak^ attaining the highest score (0.931±0.002; mean±standard deviation (std) across 10 model refits with 50% subsampled cells; Methods) and Vasc^Braak^ achieving lowest (0.698±0.017). For Abeta prediction, Ast^Abeta^ performed best (0.606±0.004), while Ex3^Abeta^, Oli^Abeta^ and Vasc^Abeta^ performances were below chance. In addition, all models significantly exceeded the 95% confidence interval (CI) of SEA-AD empirical null distribution generated with models trained on shuffled pathology labels (ROSMAP-PFC label permuted datasets, 1000 iterations; Supplementary Fig. 2a, b; Methods).

Using an alternate metric suitable for pathology class imbalance in SEA-AD (Methods), all models were further evaluated using area under the precision-recall curve (AUPRC). For Braak, all 9 models surpassed baseline AUPRC (prevalence of positive, high pathology, class). The best ΔAUPRC (model - baseline) was observed for Ex2^Braak^ at 0.509±0.010 (mean±std). Notably, Oli^Abeta^ achieved a modest improvement of ΔAUPRC=0.100±0.005. In contrast, Ex3^Abeta^ and Vasc ^Abeta^ were below baseline (ΔAUPRC<0). Here, given the systematic underperformance of all Vasc-derived models, we excluded Vasc cells from subsequent analyses.

For excitatory neurons, we wondered whether the underperformance of Ex3^Abeta^ could reflect unmodelled biological heterogeneity within this cell population, consistent with subtype specific pathological response shown elsewhere ^6,17,19^. We therefore revised the estimated pathology of SEA-AD Ex3 cells (inferred) with additional subtype-specific estimators (nine Braak and nine Abeta models trained in ROSMAP-PFC for 9 Ex3 subtypes; Methods). This refinement substantially improved predictive performance: every Ex3 subtype-specific model achieved above-chance ROC-AUC and AUPRC values (Ex3^Abeta^ results in Figure 2i; Ex3^Braak^ in Supplementary Fig. 3a-e). This improvement indicated that the initial low Ex3^Abeta^ performance deficit can be explained by biologically meaningful subtype heterogeneity rather than a general limitation of the modelling framework (see Supplementary Fig. 3f, g for results on Ex2 subtype performance).

In addition to the aggregate metrics (across model confidence thresholds; AUPRC and ROC-AUC above), we evaluated model performance as a function of score percentile thresholds (Figure 2g, h, j; see Methods). For each inferred cell type and pathology marker, the percentile threshold-dependent precision increased monotonically with higher thresholds (Vasc excluded; see above). By the 90^th^ percentile (*q*_0.9_), precisions were consistently above chance level baseline. This included Oli^Abeta^ which demonstrated clear gains at *q*_0.9_. Thus, higher predicted Braak scores were associated with increased precision, reflecting a greater proportion of true positives among high-confidence predictions. This analysis demonstrated top score prediction reliability — higher model scores reliably prioritized true positives with a proportional reduction of false positives among retained predictions. This ranking-based behaviour underpinned all downstream analyses that depended on the highest scoring subsets of cells.

Taken together, our preparatory analyses showed that ROSMAP-PFC trained Braak and Abeta estimators derived in the PFC could successfully extrapolate neuropathology signatures to another part of the human cerebral cortex—the MTG. Further, across all cell types, Braak models generalized robustly across these two cortical locations, and Abeta models exhibited substantial variability across cell types. Whether such generalizability and robustness extended to neuroanatomically much more distinct subcortical domains was the motivating question of our next analysis.

### Neocortex to non-neocortical validations of neuropathology estimators

After demonstrating PFC to MTG generalization of our five neuropathological marker estimators (cf. above), we next examined whether these generalizations persisted outside the higher association cortex, including regions outside the neocortex. For this, we utilized a multi-region dataset ^5^ (dataset referred to as ROSMAP-MR; Methods) comprised of 679,252 cells from 6 brain regions—entorhinal cortex (EC), hippocampus (HC), and thalamus (TH), in addition to PFC, midtemporal cortex (MT), and angular gyrus (AG). After inferring ROSMAP-PFC cell type identities (Supplementary Fig. 4a) and matched cell type-specific pathology-proxy scores (see above; Methods), we assessed the performance of the Braak and Abeta estimators across these diverse regions.

We evaluated cross-region model performance by examining the pathology-proxy score distributions for each donor, across the six brain regions (Fig. 3; Supplementary Fig. 4b, c). A cell type aggregated measure showed that the pathology estimates coherently tracked donor- and region-level reference pathology—donors with higher reference pathology had higher model assigned pathology scores (Figure 3a; *q*_0.9,avg_; Methods). Further, for Braak estimators, we observed strong positive correlations between known reference pathology and estimated scores, across most brain regions and cell types (maximum *τ_b_* = 0.54 for Imm^Braak^; exception: Ex2^Braak^ = −0.2 in TH; Figure 3b; Supplementary table 7). In contrast, Abeta scores displayed strong positive associations with reference pathology in PFC, MT and EC, and weak or negative associations in AG, HC, and TH, across all examined cell types (*τ_b_*< 0 in Ast ^Abeta^, Ex1^Abeta^, and Inh^Abeta^).

**Figure 3.**
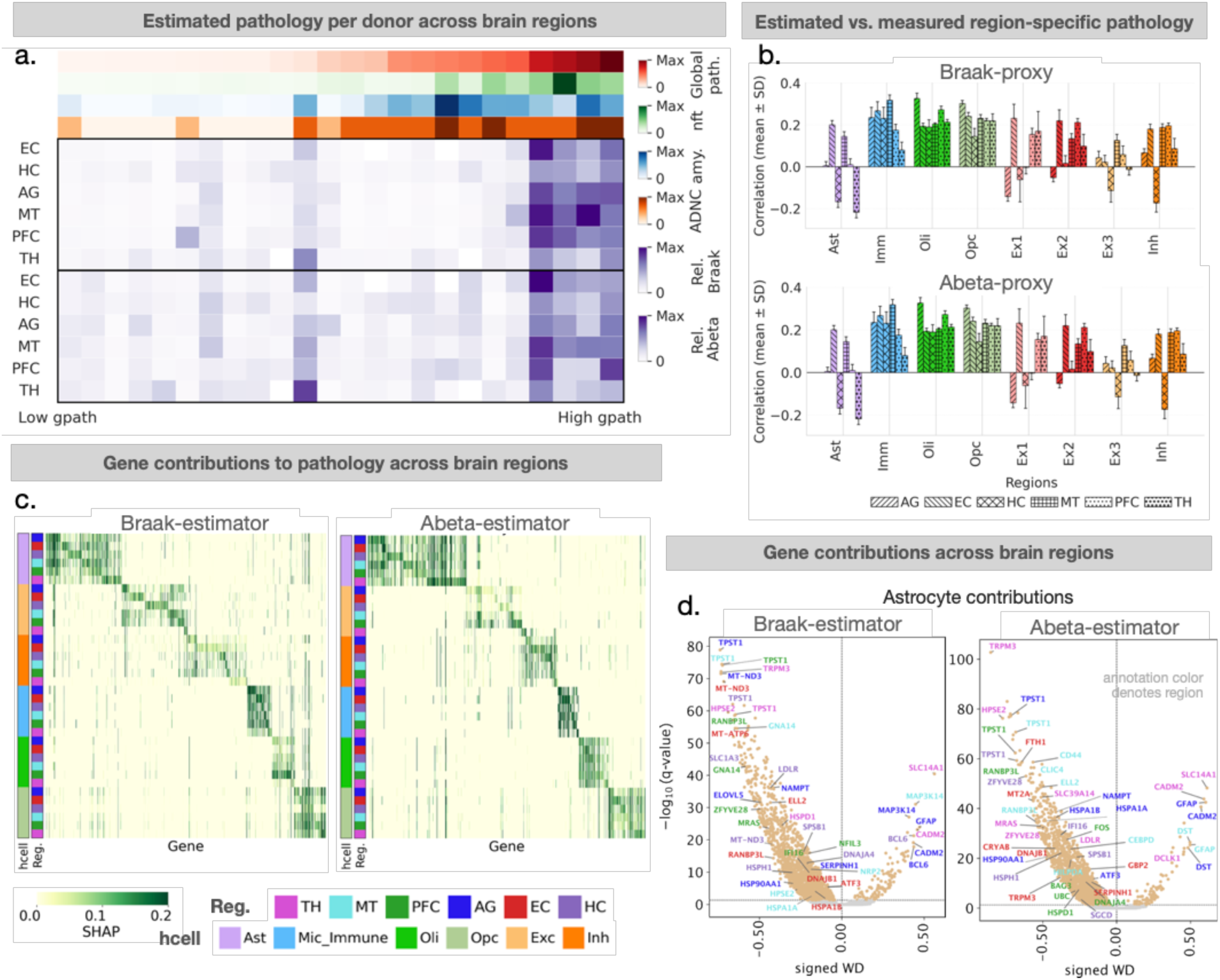
Successful transfer of neuropathology estimation models from cortical dataset to non-cortical brain regions. We validated the usability of ROSMAP-PFC trained pathology estimation models across six brain regions in ROSMAP-MR. a. Donor-level pathology across brain regions. Heatmap shows estimated pathology for 24 male donors. Each column represents a donor. The first section (four rows) shows reference pathologies (ground truth) as defined by RADC. The second section, demarcated by a bold border, represents Braak-proxy estimates. The third section represents Abeta-proxy estimates. Each row in the second and third section denotes a brain region. Colors denote 90th percentile (*q*_0.9_) of estimated pathology scores per brain region, across all cells. Darker colors represent higher scores. b. Association between reference pathology and estimated region-specific pathology load, across brain regions and cell types. Reference pathologies are region specific. Associations were quantified using Kendall’s tau-b. Bar colors represent inferred cell type; bar hatching indicates brain region. Error bars are derived from 10 model refits in ROSMAP-PFC (50% randomly subsampled cells; Methods). Top, Braak model; bottom, Abeta model. c. Gene-level contributions across brain regions and cell types. Heatmaps show differential SHAP values (*q*_0.9_ vs *q*_0.1_), computed per brain region and ROSMAP-MR cell type (Mann-Whitney U test; FDR *p*<0.05). The first colored column shows the cell type labels. The second column indicates brain region. Contribution values are normalized to [0, 1] per row. Left, Braak; right, Abeta. d. Gene contributions in Ast across brain regions. Each dot represents a gene colored by brain region. The top 20 significant genes with the highest differential SHAP (*q*_0.9_ vs *q*_0.1_) are annotated (FDR corrected *p*<0.05; Methods).

Together, these results revealed a region- and cell type-specific applicability of Abeta estimators, while supporting broad regional generalization capacity of the Braak estimators.

### Gene-level contributions to neuropathology cell signatures: going beyond cell type-level

In addition to evaluating the spatial generalizability of our neuropathology estimators, we investigated their interpretability at the gene level. Specifically, we asked whether region-level gene-effects could be recovered from our cell type-level estimators. To this end, we decomposed each ROSMAP-MR cell’s predicted pathology load into its underlying gene-level contributions using SHAP, an AI model interpretability tool (see Methods). We could then summarize these gene contributions across regions and cell types (Figure 3c, d; Supplementary Fig. 5-6; see Methods)—helping us investigate why the pathology was identified.

We examined the resulting region- and cell type-specific gene attribution profiles. Across all six ROSMAP-MR brain regions examined (see above), SHAP derived genes showed significant overlap with differentially expressed genes for both Braak and Abeta, previously identified in the same dataset ^5^ (hypergeometric test *p* < 0.05; Supplementary Fig. 4c). The extent of overlap varied by region and cell type: for Braak, the overlap ranged from 282 excitatory neuron genes in the AG to 22 Opc genes in the PFC, while for Abeta, they ranged from 141 astrocyte genes in AG to 31 inhibitory neuron genes in EC. Together, these observations demonstrated concordance between our estimator-based gene attribution framework and previously derived marker relevant genes, despite differences in the analytical approach.

To characterize regional patterns in gene-level neuropathology signals, we quantified pairwise similarity in SHAP gene-effects across the six ROSMAP-MR brain regions (see above). These comparisons revealed high concordance in pathological gene signatures between PFC and MT (adjusted *R*^2^ = 0.85 for Braak and 0.90 for Abeta, averaged across cell types; Supplementary Fig. 4d, e; see Methods). Among allocortical regions, the highest concordance was between EC and HC, as would be expected (0.76 for Braak and 0.81 for Abeta). The TH emerged as the most distinct region with the lowest concordance in pathology-associated gene contributions relative to other regions (maximum adjusted *R*^2^ was between TH-AG, 0.55 for Braak and 0.49 for Abeta). These patterns indicated shared transcriptional signatures of neuropathology among higher order cortical regions and divergent signatures between cortical-subcortical domains, consistent with the region-level relationships reported in the original study ^5^.

Collectively, these results demonstrated that our interpretability method for gene-level attribution of pathology-proxy scores recovered region- and cell type-resolved genes that were consistent with alternative gene-prioritization strategies (differential gene expression) applied to the same underlying dataset. This provided empirical support for AI interpretability-derived gene attributions for gene-level analysis of pathology-proxy across the whole-brain *Siletti* atlas (see below).

### Mapping Alzheimer’s neuropathology across the whole adult human brain

Having established the generalizability of our ROSMAP-PFC derived pathology estimators to unseen neocortical and sub-neocortical regions (cf. above), we turned to our primary goal of scientific interest: carrying over these models for annotation of the *Siletti* atlas—today’s only transcriptomic atlas of the whole adult human brain ^17^.

We applied our previously obtained and validated pathology estimators to ∼3 million cell transcriptomes from 108 brain regions, across the whole adult human brain. Following this exercise, we were in a position to derive tentative maps of Braak- and Abeta-load across finely resolved brainstem, subcortical, and cortical structures. We also derived additional maps for CERAD, TDP-43, Lewy body, as well as a negative control map for age (see below; Extended data and figures).

Prior to detailed examination of the 108 brain regions, we charted Braak and Abeta scores grouped within 10 broad brains territories — cerebral cortex, paleocortex, hippocampus, hypothalamus, thalamus, epithalamus, cerebral nuclei, midbrain, hindbrain, and spinal cord (Figure 4a-d; Methods). For each broad territory and inferred cell type, average pathology-proxy scores were low (Supplementary Fig. 7a), as would be expected. However, upon analyzing the highest vulnerability cells (*q*_0.9_ pathology-proxy score distribution percentile; Method), we observed coherent patterns of regional and cellular implications.

**Figure 4.**
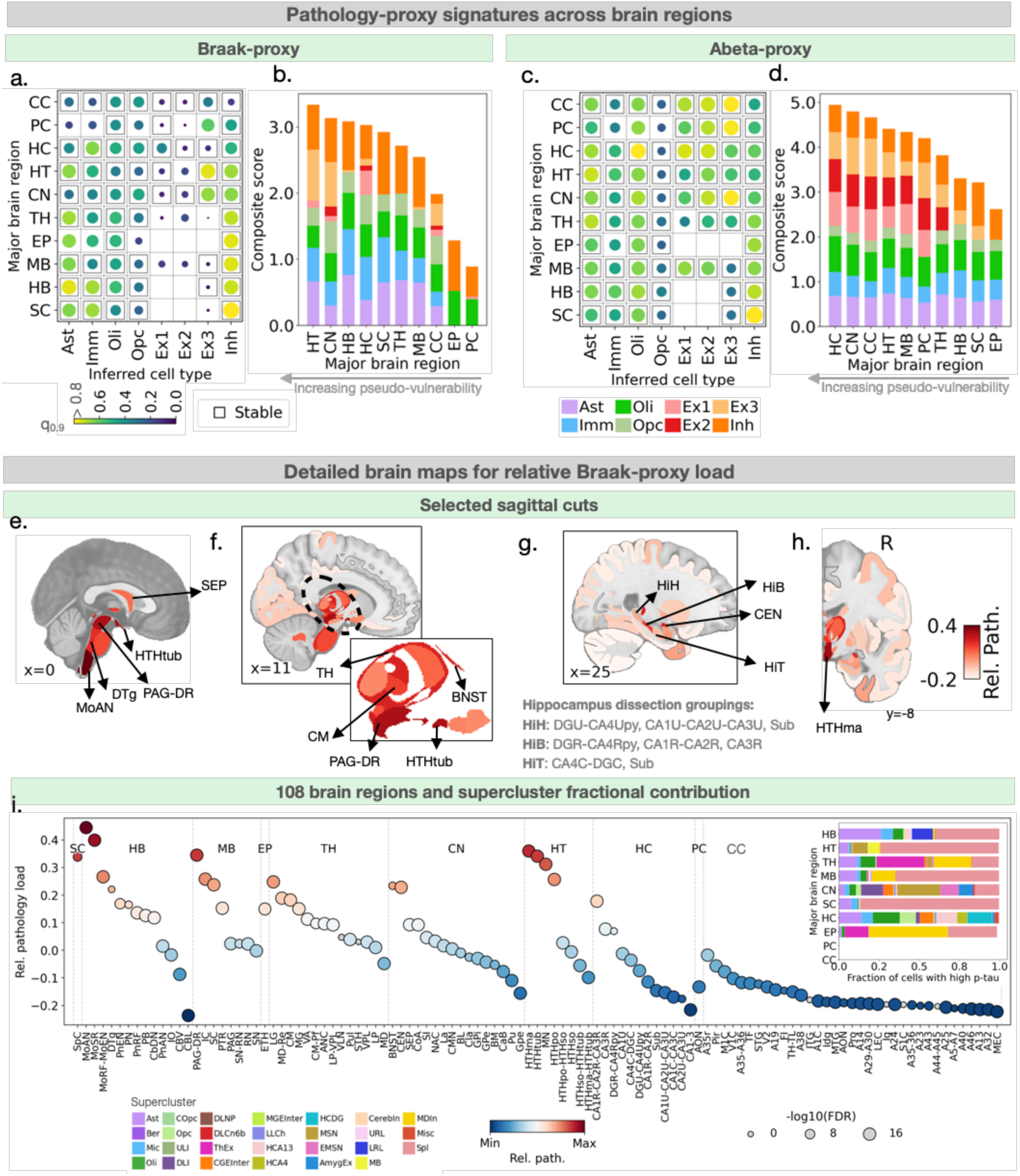
Alzheimer’s-related transcriptomic signatures manifest in distinct regions in a whole brain analysis. Estimated AD neuropathology-proxy scores across 108 brain regions in the *Siletti* snRNA-seq dataset. Braak- and Abeta-proxy scores were estimated at single-cell resolution. a-d. Aggregate summaries of estimated neuropathology-proxy score distributions across broad anatomical brain regions. a,c. Heatmap shows the 90th percentile (*q*_0.9_) of estimated pathology scores, across brain regions and ROSMAP-PFC inferred cell types. Stability was assessed using a one-sample bootstrap (n = 1000). Black outlines indicate stable estimates (c.v. of bootstrap-derived *q*_0.9_ <0.05; c.v. = standard error/mean). Empty boxes have no cells (observations). Circle size (bigger) and color (lighter/yellow) denote higher percentile scores. b,d. Contribution of ROSMAP-PFC inferred cell types across ten brain territories. Bars are stacked according to the *q*_0.9_ prediction score for each ROSMAP-PFC inferred cell type. Total bar lengths represent composite scores for a major brain region. Higher values indicate regions with stronger overall estimated pathology. e-h. Detailed maps of prominent brain areas with the highest relative Braak-proxy estimates. For each brain region, relative pathology load was computed using one-vs-rest comparison of normalized pathology estimates (z-scored within each *Siletti* defined supercluster cell population). Representative sagittal planes were selected to maximize visualization of high-scoring regions. Darker colors indicate higher relative load. Insets show enlarged views of selected sub-regions with high Braak-proxy load. i. Relative Braak-proxy estimates across 108 brain regions. Circle color and height on the plot represents the relative Braak load (one-vs-rest comparison). Circle size represents the statistical significance of the comparison (Mann-Whitney U test; FDR *q*<0.05). The inset shows the fraction of cells from a Siletti supercluster to the high scoring cells within a major brain region. All summaries were aggregated across *Siletti* donors. See Table. 1 for abbreviations.

Among 10 broad brain regions, the hippocampus emerged as top-ranking region for cell type aggregated Abeta scores and the fourth highest region for Braak scores (Figure 4b, d). Across all regions, it contributed to 12% and 11% of the total pathology-proxy burden under the Abeta and Braak models respectively. Further analysis revealed different cell types (inferred) driving the computed hippocampal vulnerability (Figure 4a). Imm cells (96% microglia) displayed the strongest Braak signal (*q*_0.9_ = 0.65; robustness assessed using a one-sampled bootstrap test with 1000 replicates), and Oli had the strongest Abeta signal (*q*_0.9_ = 0.8; Figure 4c).

Apart from the hippocampus, the Braak scores concentrated in subcortical regions (Figure 4b). The strongest vulnerabilities became apparent in the spinal cord (maximum in inferred Inh; *q*_0.9_ = 0.94), midbrain (maximum in Inh; *q*_0.9_ = 0.75), hindbrain (maximum in Ast; *q*_0.9_ = 0.73), and hypothalamus (maximum in Inh; *q*_0.9_ = 0.69). In contrast to Braak, Abeta scores were low in these regions, across cell types (Figure 4d). On the opposite end, the cerebral cortex had low Braak scores (maximum in Opc; *q*_0.9_= 0.42), but high Abeta score (maximum in Ex3; *q*_0.9_= 0.84). Thus, while Braak and Abeta scores were similar in the hippocampus, the remaining sampled brain territories diverged in their Braak- and Abeta estimates.

Across all cell types, Inh and Ast populations independently showed elevated Braak signals in several brain regions (Figure 4a). Given this observation, we examined whether the pathology-proxy signals in Inh and Ast showed concordant regional variations. We observed a strong concordance between Inh and Ast scores across 108 regions (weighted *τ_b_*= 0.42; Supplementary Fig. 8e; Methods). Among other cell populations, Oli and Opc had strong associations (weighted *τ_b_*= 0.55). These results indicated coordinated variation in pathology associated transcriptomic signatures across neuronal and glial cell populations, despite the scores being derived from separate estimators.

Together, our projected whole brain AD pathology map revealed a deep brain, subcortical emphasis of Braak and cortical emphasis of Abeta associated transcriptional signatures. This anatomical divergence in the transcriptomic gradient of Braak- and Abeta-proxies mirror some previous findings from AD post-mortem histopathology: pretangles were reported to appear earliest in the brainstem and subcortical regions, whereas plaque deposition appears in the neocortex before cascading to deeper brain structures ^10,22^. This convergence motivated a granular analysis of the Braak- and Abeta-proxy loads across 108 atlas regions.

### Braak-pathology transcriptomic vulnerability map across 108 brain regions

Given the meaningful Braak-proxy patterns observed across broad brain regions, we next analyzed these scores across 108 finely resolved *Siletti* regions (Figure 4e-i). Specifically, we mapped the relative score of each brain region compared to the rest, with higher scores denoting more vulnerability (Methods). As a control, we plotted regional vulnerability to our age-proxy (Supplementary Fig. 7c). While the age-proxy showed similar vulnerability across regions, the Braak-proxy exhibited distinct regional patterns. We visualized these anatomical patterns using a 3D reference atlas (Methods) ^23^.

Across 108 brain regions, the strongest Braak-proxy signals emerged in the periaqueductal gray and dorsal raphe nucleus, regions from the medulla oblongata (sensory relay nuclei, medullary reticular formation, and the afferent and efferent cranial nerve nuclei), multiple hypothalamic zones (tuberal zones, mamillary region, and the preoptic complex), the mediodorsal and reuniens nucleus of the thalamus, and components of the extended amygdala including the central nuclear group and the bed nucleus of stria terminalis (FDR p<0.05; Figure 4i; Supplementary table 8; see Methods for ranking metrics and robustness analysis). In contrast, the lowest scoring subcortical regions included the inferior olivary nucleus and all sampled regions of the cerebellum.

Across neo- and allocortical areas, Braak-proxy signals were highest in rostral hippocampal regions with the rostral CA1-3 subfields emerging on top, in comparison to the subiculum, the uncal and caudal regions. On the opposite end, neocortical and periallocortical regions showed uniformly low signals, with only modest relative signal in the piriform, and perirhinal (A35, A36) cortex.

To test the robustness of our findings, a similarity analysis confirmed that the region scores were highly congruent among the Siletti donors. Specifically, the pairwise cosine similarity between each donor and cross-donor aggregate scores across 108 regions was 0.63±0.05 (mean±std across donors). We also observed consistent ranking of the highest scoring regions from an alternative ranking metric that specifically accounted for changes in cell composition (Methods). This indicated that the observed regional emphasis was unlikely to be a methodological artifact.

In summary, estimating neuropathology across the whole brain, our Braak measure spotlighted discrete brainstem and subcortical nuclei, including the dorsal raphe, medullary sensory and autonomic nuclei, select hypothalamic regions, limbic-associated thalamic nuclei (mediodorsal and reuniens), and components of the extended amygdala. Within allocortical structures, the strongest signals localized to rostral CA1-3. In contrast, the cerebellum and neocortical regions exhibited low scores, suggesting Braak resilience in these regions. Overall, our analysis highlighted a highly selective and subcortical emphasis that may have been overlooked in previous AD studies.

### Cluster-specific transcriptomic vulnerability to Braak stage pathology

The human brain is extraordinarily heterogeneous in its cellular architechture: the *Siletti* atlas catalogues 31 broad superclusters and 461 finer clusters of cell popualations across the brain. To identify the specific cell populations (clusters) driving the observed Braak-proxy signals, we computed the relative contribution of each cluster to cells with high Braak-proxy scores within ten broad brain territories (Methods).

#### Non-neuronal populations

Among non-neuronal populations, astrocyte clusters showed widespread subcortical enrichment across the spinal cord, midbrain, hindbrain, thalamus, and hypothalamus (Figure 4a; Supplementary table 9). Differential SHAP analysis on gene roles identified APOE (lipid transport driver), CST3 (protease inhibitor), SLC1A2 (glutamate clearance), and CD44 (disease associated astrocytes) as major contributors to elevated pathology scores across these regions (FDR *q* < 0.05; Figure 5; Supplementary table 11; Methods). Indeed, prior transcriptomic studies that have shown alterations in these genes are characteristic of reactive, disease-associated astrocytic states in neurodegeneration ^24^.

**Figure 5.**
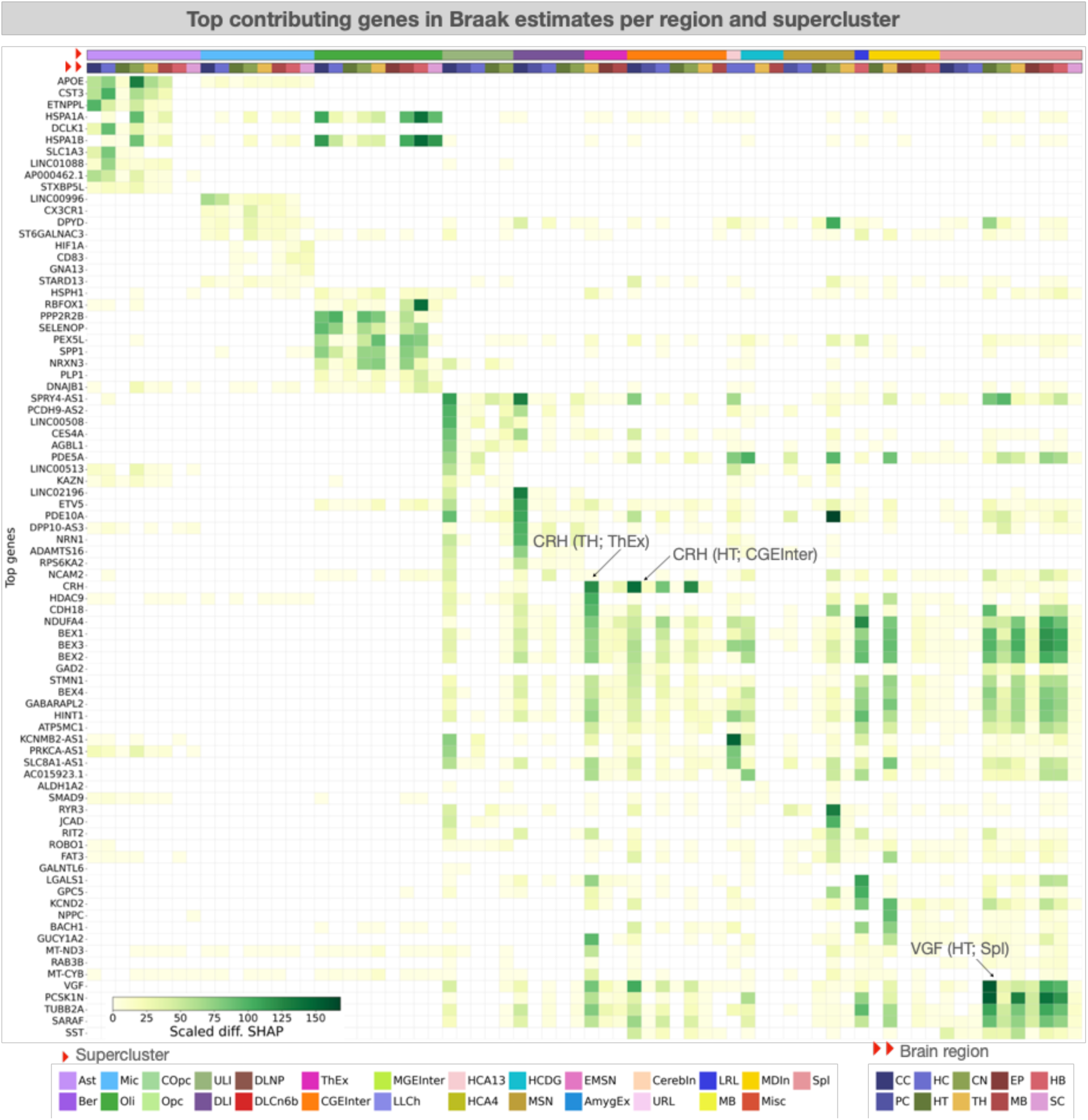
Key genes contributing to Braak-proxy vulnerability in major cell populations, across brain regions. Heatmap shows the contribution of genes to estimated Braak-proxy (top 3 per supercluster and brain region). Color intensity indicates effect size (q <0.05; Scaled differential (diff.) SHAP = √diff. SHAP*-log10(q); Methods). Illustration done for select superclusters with high Braak-proxy estimates. Annotations with arrows depict notable genes with corresponding region and supercluster in brackets.

**Figure 6.**
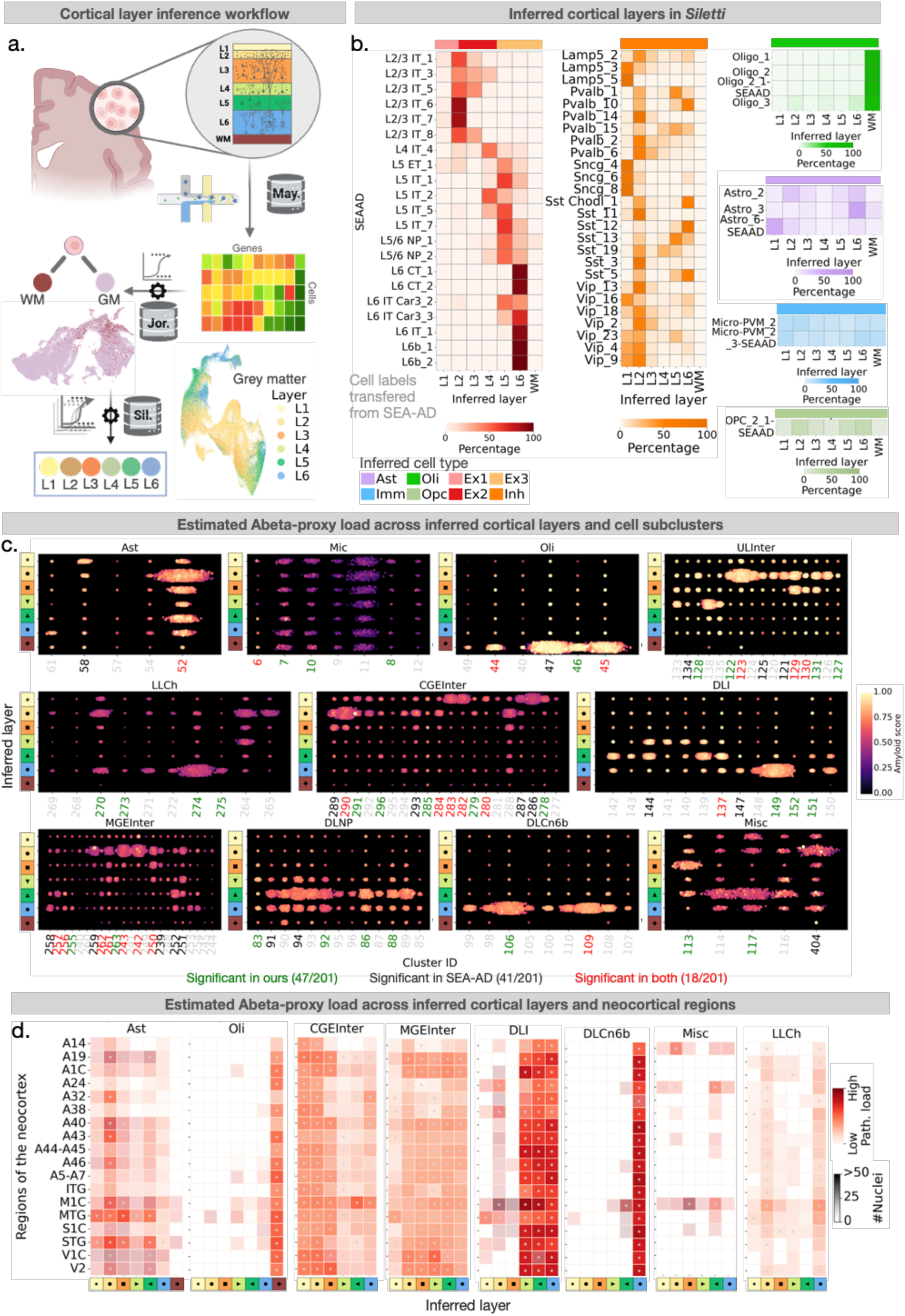
β-Amyloid related transcriptomic signatures show laminar organization across the neocortex. a. Schematic of single cell layer classifier. First, a binary classifier, trained on *Maynard* snRNA-seq dataset, distinguished white matter (WM) from grey matter (GM) cells. Next, for the cells classified as GM, a multi-class logistic regression model trained on *Jorstad* snRNA-seq dataset was used to assign cells to one of six neocortical layers (layers L1 - L6). Figure created using BioRender. b. Inferred layers for neocortical cells in the *Siletti* dataset. For illustration, cell annotations were transferred from Human MTG SEA-AD using *MapMyCells* ^34^. Grid color intensity indicates the percentage of cells of a given type within each inferred layer. Inferred layers align with expected laminar origin; for example, L6 CT neurons localize predominantly to L6, L2/3 IT neurons distribute across L2 and L3. c. Distributions of estimated Abeta-proxy load across laminar layers and superclusters. Subplots represent *Siletti* defined superclusters. Each dot represents a cell colored by inferred pathology score (lighter color represents higher score). Cells are stratified by model inferred cortical layer (y-axis) and *Siletti* defined subcluster (x-axis). Pseudo-vulnerable cell subclusters were identified by comparing pathology scores within each subcluster to scores of the remaining cells (Mann-Whitney U test; FDR corrected *q*<0.05). Pseudo-vulnerable subclusters are annotated in red if implicated as vulnerable elsewhere ^19^. d. Estimated Abeta-proxy load across inferred laminae and brain region. Each heatmap grid shows the 90th percentile (*q*_0.9_) of estimated pathology scores across neocortical subregions. Color intensity reflects *q*_0.9_ scores (darker indicates higher value). The transparency of a grid represents the number of cells exceeding the *q*_0.9_ threshold within each supercluster. Stability of *q*_0.9_ estimates was assessed using bootstrap resampling (n=1000 iterations). Stable region-layer combinations (coefficient of variation < 0.05) are marked with grey stars.

In contrast, microglia populations exhibited significant enrichment among high Braak cells only in the spinal cord, hindbrain, and hippocampus (Figure 4a). Pathology-associated genes converged on immune and regulatory programs. Prominent contributors included APOE (associated with lipid remodelling), HLA-DRA (antigen presentation), FOXP1 (transcriptional control), DPYD (metabolic adaptation), and ASAP1 (cytoskeletal reorganization) — consistent with immune-activated, disease-associated microglial states ^25^.

#### Subcortical, deep brain neuronal populations

Among neuronal populations, splatter neurons (Spl) expressing VGLUT2 were consistently enriched among high Braak scoring cells from the spinal cord, midbrain, hindbrain, and hypothalamus. Specifically, VGLUT2^+^ Spl neurons contributed 53% of all high scoring cells (HSCs) in the spinal cord (despite comprising 36% of the overall Spl population in the spinal cord), 40% in the midbrain (37% Spl^VGLUT2^ overall), 30% in the hypothalamus (28% Spl^VGLUT2^ overall). This preferential enrichment of VGLUT2^+^ Spl was notable given the high heterogeneity of the Spl population (22% of all Spl neurons were GABAergic, 39% glutamatergic, 1.4% co-expressed both, as well as smaller cholinergic, glycinergic, serotonergic and dopaminergic subpopulations ^17^).

Elevated pathology-proxy scores in Spl neurons were driven by differential SHAP genes including PCSK1N, SST, VGF, SCG2, GNAS (neuropeptide processing and secretory signaling), BEX1/2/3, CDH18 (synaptic adhesion and neuronal maintenance), TUBB2A (microtubule integrity), and NDUFA4 (mitochondrial respiration). Collectively, vulnerable VGLUT2+ Spl neurons implicated genes related to synaptic vulnerability and disruptions in metabolic activity (Figure 5a).

We next investigated the thalamus. Here, thalamic excitatory neurons expressing VGLUT2 contributed the highest proportion of HSCs (29%; 14% of all thalamic cells). The top driving gene in this population was CRH, which encodes the corticotropin-releasing hormone, a hallmark regulator of stress-axis signaling. Alterations to CRH mRNA levels have been previously reported in AD ^26^. In addition, we observed significant contributions from the PVALB gene. Although PVALB has been linked to AD-vulnerable inhibitory interneurons in the cortex ^5,19^, its role within excitatory neuron populations remains unknown.

Given the elevated Braak estimates in the medulla oblongata — a region not typically emphasized in AD — we further examined the cellular sources of these signals. Within the sensory relay nuclei, Spl^VGLUT2^ neurons accounted for the largest fraction of high-scoring cells (15% of all HSCs from sensory relay nuclei). In contrast, within the cranial nerve nuclei, the signal was distributed across oligodendrocytes (14% HSCs from cranial nerve nuclei), microglia (8%), and Spl^VGLUT2^ neurons (8%).

The identity of the highest Braak-enriched cell class shifted progressively across examined brain regions — transitioning from VGLUT2+ neurons (see above) to mixed VGLUT1/2 populations in the hindbrain, and VGLUT1+ in the hippocampus. Across all sub-regions within the hindbrain, Spl^VGLUT2^ contributed 20% of the overall HSCs from hindbrain, followed by lower rhombic lip neurons expressing VGLUT1/2 (13% of HSCs; top SHAP genes included NDUFA4, LGALS1 and GPC5).

Overall, within the brainstem, selective neuronal populations expressing VGLUT1/2 dominated the high Braak signatures bearing cell populations.

#### Hippocampal neuronal populations

Within the hippocampus, the strongest Braak signals emerged in granule cells (15% of hippocampal HSCs; multiple ribosomal RPS/L genes) and CA1 pyramidal neurons (13% of hippocampal HSCs; top SHAP genes: NNAT, multiple antisense transcripts including KCNMB2, PRKCA, SLC8A1), both expressing VGLUT1. This pattern was consistent with prior histological and transcriptomic-epigenomic studies identifying hippocampal granule cells and CA1 pyramidal neurons as particularly vulnerable, suffering early epigenomic information loss associated with cognitive decline in AD ^5,6,22^.

In contrast to the high transcriptomic vulnerability of the glutamatergic neuron populations from the subcortical regions, the lowest Braak-proxy contributors included GABAergic neurons (despite raw proportions of GABAergic neurons ranging between 66% of all neurons in the thalamus to 15% in the spinal cord), oligodendrocyte precursor cells, neocortical neuron populations (e.g., the deep layer near projecting neurons), mammillary body neurons, and the upper rhombic lip neurons projecting to the cerebellum.

To anchor cell type-resolved Braak transcriptional signatures to an independent, clinically accessible biomarker modality, we also compared the SHAP genes across cell types against early Braak-associated cerebrospinal fluid (CSF) proteins. These were identified in a previously published study (preclinical Aβ+tau+ versus control) ^27^. A total of 133 SHAP genes from our study significantly overlapped with these early CSF markers (137/323 implicated proteins; Supplementary table 12; hypergeometric test *p* = 2.5×10^−9^; see Methods). Among these, 64 SHAP-CSF genes were associated with splatter neurons (VGF, SCG2, and NEGR1; top 3), 39 with astrocytes (LUZP2, APOE, and CST3), and 11 with microglia (APOE, HSPA1A, and SPON1). These findings nominate the likely cellular sources of CSF molecular markers detectable during the preclinical stages of AD.

Here we delineated region specific cell populations expressing high Braak-associated transcriptional signatures in a brain-wide context. In summary, astrocytes demonstrated wide-spread brainstem and subcortical elevation of Braak-proxy signals. Microglia showed the high signals within the hippocampus. Among neuronal populations, selective VGLUT1/2+ subpopulations exhibited elevated Braak-proxy signals. In contrast, GABAergic populations showed consistently low Braak-proxy scores.

### Transcriptomic signatures of β-amyloid plaque across brain regions, cell types and inferred cortical laminae

We next analyzed the microanatomical and cellular distributions of estimated Abeta-proxy load in the *Siletti* atlas. As shown above (Figure 4d), cell type aggregated Abeta-proxy signals were highest in the cortical regions (neocortex, periallocortex and allocortex). Peak Abeta-proxy load localized to the rostral hippocampus, where the CA1-CA3 subfields showed the highest scores across all cortical regions (Supplementary Fig. 7b). The primary contributions were from hippocampal granule cells and CA1 pyramidal neurons (Supplementary table 11). Notably, this pattern mirrored the Braak-proxy signals within the hippocampus (Figure 4i), suggesting these hippocampal regions and cell populations had similar transcriptomic vulnerability to Braak and Abeta proxies.

Moreover, we previously described a spatial divergence between Braak- and Abeta-proxy distributions across the broader brain territories (subcortical emphasis for Braak and cortical for Abeta; see above). This dissociation persisted within the cortex at finer resolutions. Allocortical regions, including piriform, and perirhinal cortices, displayed the lowest Abeta-proxy scores (in contrast to high Braak-proxy signals), whereas the neocortical regions showed comparatively higher burden.

Within the neocortex, the highest relative signals were observed in the primary motor cortex (significant at FDR *q* < 0.05; Methods), followed by the primary and secondary visual cortices, primary auditory cortex, and the primary somatosensory cortex (Supplementary Fig. 7b). However, differences in the relative scores across these regions was minor, indicating a broadly uniform transcriptomic Abeta signature. This pattern is consistent with immunochemical and PET studies of early AD reporting relatively homogeneous distribution of pathological plaque across neocortical areas ^28–30^.

Laminar differences in vulnerability to AD neuropathology have been described in prior studies of advanced AD ^19,31^. We therefore asked whether such layer-specificity could be detected in our transcriptomic whole brain model of Abeta. To this end, we inferred neocortical layer identities for cells in the *Siletti* atlas using a secondary classifier developed using cortical lamina specific snRNA-seq datasets (*Jorstad* ^32^*, and Maynard* ^33^; Methods). This added another kind of critical information to *Siletti* which was not provided as part of its own meta data (**Error! Reference source not found.**a, b).

Across neocortical layers, the strongest Abeta-proxy signatures emerged in layers L2 and L5-L6. In contrast, the weakest signals were observed in L1 and L4 (**Error! Reference source not found.**c, d; Supplementary table 13). These layer-specificities were attributable to distinct pseudo-vulnerable neuronal and glial populations (FDR *q* < 0.05; see Methods). In the upper layers (L2-L3), L2/3 intratelencephalic (IT) neurons showed the highest Abeta-proxy load (top genes included AGBL1, PDE10A, HOMER1; Supplementary table 14), along with somatostatin+ medial ganglionic eminence neurons in L3 (SST, VGF), fitting with previously reported AD-vulnerability of these cell populations ^19^. In deeper layers (L5-L6), the highly active, long-range projecting IT and corticothalamic neurons (DPP10-AS3, SPRY4-AS1, ETV5) showed elevated Abeta-proxy loads, compared to the weaker signals in low-activity L6b neurons.

In addition to layer specific neuronal contributions, pseudo-vulnerable glial cells preferentially localized to specific cortical layers (Supplementary table 13). Pseudo-vulnerable astrocytes were primarily localized to L2 (35% of all pseudo-vulnerable astrocytes), and the rest were distributed across L3 (19%) and L5 (18%) — broadly mirroring the laminar distributions of pseudo-vulnerable neuronal populations (see above). This astrocytic laminar pattern is consistent with recent work showing that AD-associated astrocyte transcriptional substates are non-uniformly distributed across the cortical depth with preferential enrichment in upper cortical layers for specific reactive clusters ^31^. We further observed that pseudo-vulnerable microglia were concentrated in L1(21%) and the white matter (20%), suggesting a similar layer-specific enrichment of pathology-associated microglia.

In summary, across the cortex, Abeta-proxy signals were strongest in the neocortex compared to allocortical regions. The neocortical signals were broadly uniform across fine-grained areas. In contrast, pronounced laminar selectivity was observed, with enrichment of Abeta-proxy signals in L2 and L5-L6 which was driven by distinct cell populations: L2/3 IT neurons drove upper layer signals and the deep-layer IT and corticothalamic neurons drove the deeper layer enrichment. Glial cells also exhibited layer-specific patterns, with pseudo-vulnerable astrocytes primarily enriched in L2 and microglia in L1 and the white matter.

### Regulatory mechanisms selectively driving high pathology signatures

In addition to mapping AD pathology signatures across the whole human brain, we sought to identify the biological mechanisms associated with cellular and regional vulnerability. To this end, we first used an independent analysis to define the transcriptional regulatory programs active in each cell (SCENIC; Methods). We next tested which of these programs were preferentially active in cells with the highest estimated pathology burden, thereby identifying candidate regulatory mechanisms associated with vulnerable cellular states (FDR *q* < 0.05; Methods).

Across all brain regions and superclusters, we identified 424 regulons (transcription factors and their target gene sets) that were significantly associated with pathology-proxy signatures of either Braak or Abeta (2308 active regulons across the whole brain). Out of these, the highest number were in microglia (47/424), astrocytes (45/424) and splatter neurons (42/424) (a; Supplementary table 15-16). This was consistent with the observed high transcriptomic vulnerability of these cell populations (see above).

Within astrocytes, the top Braak-proxy-associated regulons were enriched for functional processes related to synaptic plasticity/memory, MAPK signaling, redox metal ions, oxidative stress and apoptosis (CEBPD(+), JUNB(+), ATF3(+)), calcium signalling and glucose metabolism (FOS(+)), and angiogenesis and cytoskeletal structure (BCL6(+)) (across GO, KEGG and SynGO pathway databases; b, d; see Methods). In contrast, microglia associated regulons were related to inflammatory signals (REL(+), CREB1(+)), viral processes (FOS(+), JUNB(+)), protein ubiquitination (ETV6(+), FLI1(+)), and lipid/glucose metabolism (including HIF1A(+)). These enrichments are consistent with previously reported AD-associated astrocytic and microglial dysfunctions ^7,31,35,36^, corroborating the pathological relevance of our model-derived pseudo-vulnerable cell populations.

Within neuronal cell populations, regulons associated with high Braak-proxy scores were enriched for terms related to synaptic signaling and transmission (FOS(+)), neurogenesis (CUX1(+)), cell adhesion (JUND(+)), and cytoskeletal structure regulation (PBX1(+)). Oligodendrocyte lineage cells with the highest neuropathology signatures were associated with synaptic structure and maintenance (HIVEP2(+)), myelination (SOX10(+)) and lipid metabolism (PAX3(+)).

Further, transcription factors regulating circadian rhythm showed involvement in almost all pseudo-vulnerable cell populations (from both Braak and Abeta; Figure 7d). Specifically, these TFs regulated the CEBPD(+) regulon in astrocytes and microglia, SREBF2(+) in splatter neurons and upper layer IT neurons, ZBTB7C(+) in thalamic excitatory neurons, and THRB(+) in L2/3 IT neurons, among others (c, d). This is consistent with recent findings from human epigenomics ^6^ and a mouse translatome analysis showing reprogramming of circadian rhythm gene expressions in plaque affected cells^37^.

**Figure 7.**
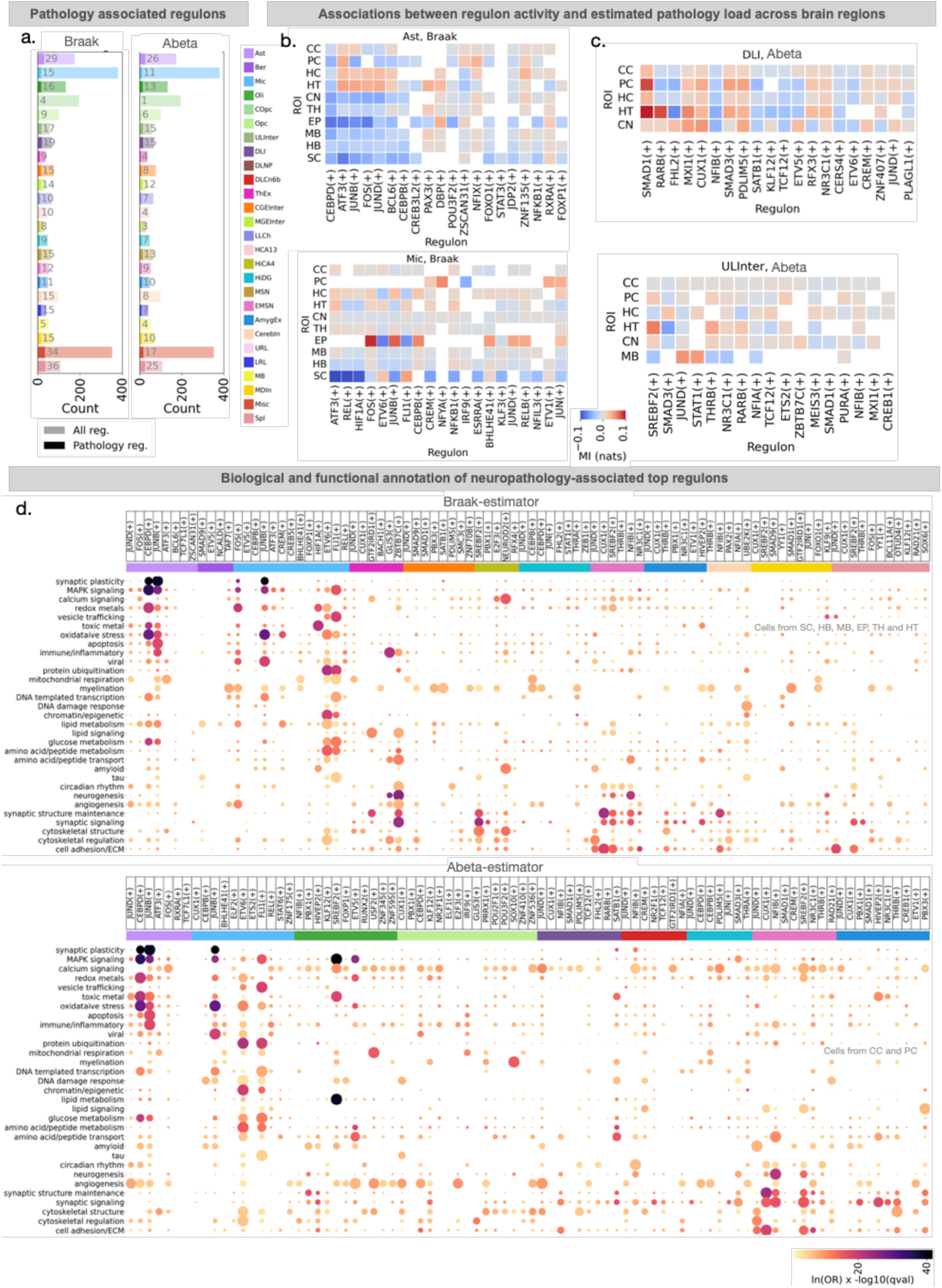
Whole-brain distribution of AD-associated regulatory mechanisms. a. Proportion of pathology associated regulons among all detected regulons. Regulons are defined as groups of transcription factors and their target genes (SCENIC). Light bars indicate the total number of regulons per supercluster across regions; dark bars indicate regulons significantly associated with estimated pathology. Associations were defined by mutual information (MI) between regulon activity and pathology scores within each supercluster. Significance was assessed by permutation of regulon activity scores (n=1000 iterations; observed value outside two-sided 90% confidence interval). Bars are colored by supercluster. b-c. Regional and supercluster specificity of regulon-pathology associations. Heatmap shows the top 20 pathology-associated regulons ranked by strength of association with estimated pathology. Color intensity represents MI; direction of association was assessed using Kendall’s tau-b (dark red indicates strong positive association. Dark blue indicates strong negative association). Significant associations in selected superclusters are shown (Left, Braak; right, Abeta). d. Gene set enrichment analysis of pathology-associated regulons. (Top) Braak enriched terms in SC, HB, and MB. (Bottom) Abeta enriched terms in PC and CC. Terms were grouped into biological topics by semantic filtering. Each dot represents a term. Dot color represents the *Enrichr* log(odds ratio), scaled by -log10(FDR corrected *q*-value). Top colored bars indicate supercluster; Top column annotations mark regulons enriched for the indicated topic.

Together, these analyses revealed brain region- and cell type-specificity of the regulatory landscape underlying AD associated neuropathology proxies. Broadly, prominent gene programs were linked to synaptic plasticity and memory, apoptosis, oxidative stress, circadian rhythm, and neurogenesis, with region and cell type dependent up- or down-regulation.

## Discussion

Despite the transformative impact of single-cell genomics, its application in disease across the entire human brain remains limited by cost, logistics, and technical feasibility. This constraint has impeded our understanding of the molecular landscape of AD to a limited set of cortical regions thus far submitted to patient-level snRNA-seq profiling. Yet evidence from human brain autopsy consistently suggests involvement of subcortical brain regions in the earliest disease stages. Here, we demonstrate a path towards a *whole brain,* cell type resolved atlas of Alzheimer’s neuropathology inferred from transcriptomics. To achieve this goal, we devised a data-driven protocol to extend AD-critical molecular signals to single cells in the recently available *Siletti* resource — the largest brain transcriptomic resource in healthy humans, with ∼3 million cells. By deriving Braak stage and β-amyloid annotations for each *Siletti* cell, we could map pathology-associated molecular similarity across brain regions, cell types, and inferred neocortical layers.

Tailoring a computational framework for this purpose, we derived cell type-specific transcriptomic proxies of neuropathologies from the ROSMAP-PFC resource. These proxies capture detectable signatures of molecular processes associated with pathology burden, rather than directly observed pathology-lesions. By means of the devised machine learning protocol, we could perform the inference of disease-associated states for every separate cell at hand. Notably, we showed that these signals in cell transcriptomes are distinct from general aging-related signatures, reinforcing the view that the uncovered transcriptomic signatures reflect neuropathology-specific molecular alterations. Validations in independent AD datasets further demonstrates that our derived pathology models generalize to unseen cell populations, unseen participants, and unseen brain regions.

Overall, several established insights from previous AD studies are recapitulated in our brain-wide map of pathology vulnerability. 1) The anatomical patterns of transcriptomic AD-associations we observe are not random, but replicate progression patterns established by seminal AD-staging frameworks. In particular, our Braak estimates track the expected sequence of Braak stages ^9^. 2) The biological pathways implicated by functional annotation of our model-derived genes highlight synaptic plasticity (downregulated in astrocytes and microglia), synaptic signaling, cytoskeletal structure (neuron-specific), and myelination (upregulated in oligodendrocyte precursors). These nominated pathways have been associated previously in AD transcriptomic studies of humans ^7,19,21,38,39^. 3) The structured neocortical laminar organization observed in our Abeta-burden map is consistent with prior genomic and structural studies in AD ^28,19^. Taken together, these observations support the biological coherence of our data fusion approach that extrapolates AD-related molecular signatures to a general-purpose whole brain resource.

Beyond just replicating evidence of earlier research, our whole brain AD characterization exposes several novel findings in uncharted territories. Across transcriptomic analyses, we highlight specific subcortical, deep brain cells that exhibit molecular profiles closely resembling Braak-associated cell states. Notably, several of these regions have prior histopathological evidence of early tauopathy ^40^. For example, postmortem histopathology studies have reported the dorsal raphe nucleus as one of the earliest pretangle bearing regions in middle aged individuals without dementia ^11,41,42^. Similarly, pretangle pathology has been identified in the reuniens nucleus of the thalamus ^43^, and in hypothalamic nuclei (tuberomammillary nucleus and the ventromedial hypothalamus) ^44^, before observable cortical tangles. In contrast, the cerebellum — where we observed minimal Braak-proxy signals — has indeed been reported to be relatively resilient to tau tangles ^45,46^. Thus, despite lacking disease data, we speculate that the molecular landscape of select subcortical brain regions may predispose them to subsequent pathology characteristic of AD.

The subcortical regions most vulnerable to Braak-proxy in our analysis — including the dorsal raphe, select hypothalamic zones, dorsal tegmentum (anatomically close to locus coeruleus), and the medulla oblongata — point to circadian, central autonomic, and neuroendocrine circuits as candidates of early molecular vulnerability in tauopathy. The involvement of these regions in our estimated whole brain map aligns with several longitudinal cohort studies showing sleep-wake and autonomic system disturbances in preclinical AD ^47–54^. Our findings further support investigators proposing early susceptibility of locus coeruleus-hypothalamic networks ^42,55,56^. Moreover, the cellular enrichment of circadian rhythm-related pathways across multiple populations (from our regulatory network screenings of high pathology cells) suggests circadian disruption manifests not only in vulnerable brainstem nuclei but may reflect a broader regulatory feature observable across the brain.

The single-cell resolution of the *Siletti* resource allowed us to test whether AD-like signals reflect general tissue signatures or specific cellular vulnerability. Consistent with previously proposed selective neuronal vulnerability in AD ^57^, we identified discrete cell populations with strong disease-associated signatures. Within the hippocampus, for instance, granule and pyramidal neurons from the rostral CA subfields exhibited the strongest AD signatures. These neuron subtypes have been repeatedly implicated in previous AD postmortem and omics studies ^5,57,58^, and selective vulnerability of the rostral hippocampus neurons have been previously documented ^59^. Their detection in neurotypical brains permits the assumption that the hippocampus harbors cell type-specific intrinsic molecular programs that may be conducive to aspects of AD pathology.

An analogous pattern of selective neuronal vulnerability emerged in subcortical regions. Here, glutamatergic neurons exhibited the strongest Braak-associated molecular profiles, including splatter neuron subtypes, thalamic excitatory neurons, and lower rhombic lip neurons. This pattern suggests that VGLUT1/2-expressing neurons, which maintain high baseline cytoskeletal turnover and operate near cellular stress limits, may be predisposed to tauopathy ^60–63^. The presence of disease-associated astrocytes and microglia in these anatomical compartments is consistent with reactive or compensatory responses that may accompany or modulate neuronal vulnerability. Notably, molecular vulnerability does not imply early degeneration, as seen in locus coeruleus neurons, which exhibit resilience despite substantial pathology ^13^. The cell subpopulations identified here open entry gates to study such resilience.

While we found Braak vulnerable neuronal subtypes to vary across brain regions, within the thalamus, vulnerability was particularly evident in corticotropin-releasing hormone (CRH) expressing excitatory neurons. Among these CRH subtypes, a subset also expressed PVALB, a neuromodulator usually associated with fast-spiking inhibitory neurons ^64^. Given that hyperactivity of PVALB interneurons is known to be pronounced in the initial stages of AD ^65^, it will be informative to examine whether CHR+/PVALB+ thalamic neurons constitute a distinct excitatory counterpart.

Relative to the subcortex, our analyses of the neocortex showed limited AD-likeness, with low Braak signatures and a diffuse Abeta profile. However, there was a robust laminar component to the Abeta-proxy signatures. In particular, L2-L3 neurons expressing somatostatin+ and L5-L6 long-projecting intratelencephalic neurons displayed the most prominent Abeta-proxy signatures. This is consistent with layer- and cell type-specific findings from the few transcriptomic studies of selected regions of the AD neocortex ^6,19^.

The observed neocortical laminar selectivity of Abeta-proxy signatures extended to glial populations. Our approach localized pathology-associated astrocytes to L2 and L5, and microglia especially to L1 and L6. Laminar preference of AD-associated astrocytes has been reported in a recent AD spatial-transcriptomic work ^7^, but not in microglia. Our findings add new bricks of evidence that glial contributions to AD-associated molecular states may be structured by cortical layers and warrant further investigation.

More broadly, the advent of our data-augmentation framework now motivates several therapeutic, diagnostic and exploratory applications. First, the region and cell type-specific genes and transcriptional regulators identified here provide a resource for early-stage precision medicine therapeutics — when interventions are most likely to prevent or delay the onset of symptoms. Second, by anchoring ‘usual suspect’ CSF protein markers to specific cell types (e.g., VGF from glutamatergic neurons, CST3 from astrocytes, and BASP1 from microglia), our study provides plausible interpretations of the cellular origins for these preclinical AD biomarkers ^66^. In addition, extending this framework across disorders can enable systematic comparisons of molecular, regional and cellular signatures of pathology. For example, our preliminary analysis of Lewy body-proxy markers indicates a prominent amygdaloid component, consistent with brain-first models of Parkinson’s disease ^67^ (see Supplementary Fig. 8b).

In summary, by unlocking the first anticipated whole brain map of AD, our collective findings nominate new brain regions with putative roles in AD pathology. This includes candidate regions that could not be examined in prior transcriptomic studies in humans. We further delineate the spatial, cellular, and neocortical laminar organization that may be particularly relevant in early AD. Our collective findings suggest that the molecular fingerprint of AD extends to subcortical brain regions linked to mood, sleep-wake regulation, and central autonomic functions, urging their systematic inclusion in future AD single-cell studies.

## Methods

### Datasets

ROSMAP-PFC ^21^: The snRNA-seq dataset was derived from postmortem brain tissue collected from 427 participants in the Religious Orders Study and Rush Memory and Aging Project (ROSMAP), two longitudinal clinical–pathological studies of ageing and dementia in which all participants are brain donors. The study sampled the prefrontal cortex of 212 males and 215 females. Based on modified NIA-Reagan criteria, 238 individuals had pathological AD, and 189 did not. Reads were aligned to the GRCh38 human reference genome using Cell Ranger v3.0.2, with pre-mRNA reads included to capture unspliced nuclear transcripts. Following quality control and doublet removal, the final dataset comprised 2,359,994 nuclei. Major cell-class and higher-resolution subtype annotations provided by the original authors were retained without modification.

SEA-AD ^19^: Middle temporal gyrus brain specimens were derived from donors in the Adult Changes in Thought Study and the University of Washington’s Alzheimer’s Disease Research Center. The study included participants from age groups ranging from <65 years to > 90 years. Out of 89 donors, 36 were males, and 42 subjects had recorded dementia. The mean age of the individuals was 88 years. In total we analyzed 1,230,092 cells in this dataset. In total, and 36,517 genes were recorded and mapped to hg38 (GRCh38-2020-A) human reference genome. In our study, 21 major cell types from the atlas was used for evaluation.

ROSMAP-MR ^5^: The snRNA-seq dataset was derived from postmortem brain tissue collected from 48 participants in the Religious Orders Study and the Rush Memory and Aging Project (ROSMAP), two longitudinal clinical–pathological studies of ageing and dementia in which all participants are brain donors. Individuals were selected based on modified NIA–Reagan criteria and Braak stage, including 26 participants with a pathological diagnosis of AD and 22 without AD. The groups were balanced by sex and matched for age and postmortem interval, with median ages of 86.6 years in the AD group and 86 years in the non-AD group. After quality control, doublet removal, and exclusion of clusters showing strong donor-specific batch effects, the final dataset comprised approximately 1.35 million nuclei from 283 samples. Cell identities annotated by the authors were reused. Sequencing reads were aligned to the GRCh38 human reference genome.

Siletti ^17^: The whole brain cell atlas for adult humans comprises 3.369 million nuclei from three postmortem adult donors, sampled across 108 anatomical locations. Following quality control, transcriptomic profiles were organized into a hierarchical taxonomy of 31 superclusters and 461 clusters. Throughout this study, we retained the cell-type and anatomical nomenclature defined in the original study. In total, this study recorded 59,480 genes mapped to hg38 (GRCh38. p13 gencode V35 primary sequence assembly) of the human reference genome. In this study, for X-Y duplicate genes, we retained the X-chromosomal gene.

Jorstad ^32^: The snRNA-seq dataset was generated from postmortem brain tissue obtained from five neurologically and neuropsychiatrically healthy adult donors. These included three males and two females, with a mean age of 47 years and a mean postmortem interval of 12.8 hours. Tissue was sampled from multiple cortical regions, including dorsolateral prefrontal, anterior cingulate, primary auditory, middle temporal, primary motor, primary somatosensory, primary visual, and angular cortices. In total, 49,417 nuclei were isolated from either full cortical-depth dissections or layer-specific microdissections and profiled using SMART-seq v4 or the 10x Chromium Single-Cell 3′ platform. Sequencing reads were aligned to the GRCh38 human reference genome, with both exonic and intronic reads included in gene-expression quantification to capture unspliced nuclear transcripts.

### Data preprocessing

Our study built on the processing and quality control that had already been conducted by the original authors of the eligible studies, with minimal additional preprocessing of the snRNA-seq datasets. For the training datasets, ROSMAP-PFC and *Jorstad*, we excluded cells expressing fewer than 300 different genes. Following established best practices, we applied cell-level normalization in all datasets ^68^. That is, we scaled total counts per cell to 10,000 unique molecular identifiers with scanpy *pp.normalize_total* (v1.11.2).

### Inference models’ overview

In our pathology inference pipeline (Fig. 1), multiple stages required dedicated modelling components, including cell type inference, neuropathology estimation, and neocortical layer inference. Within the context of this study, we prioritized transparency, scalability, and robustness to dataset shift (e.g., donor, batch, and platform differences), rather than models maximizing predictive accuracy ^69^. Accordingly, we used l2-regularized logistic regression models, which provide well-posed estimators in the high-dimensional setting, such as snRNA-seq datasets ^70^. In comparison, deep learning models, while powerful in capturing higher-order interactions, are less often directly interpretable at the gene level ^71,72^. In addition, compared to other classical machine learning methods such as support vector machines or gradient boosted trees, logistic regression models provide inherently probabilistic outputs, reducing the need for post hoc calibration adjustments in our investigation.

### Pathology inference model

We aimed to annotate single-cells, in suitable snRNA-seq datasets, with a pathology-likeness score based on their transcriptional similarity to cells derived from donors with established neuropathology. To this end, we developed pathology inference models to quantify transcriptomic signatures indicative of cell states related to pathology states at single-cell resolution using donor-level neuropathological annotations from the ROSMAP-PFC dataset.

In constructing the models, we considered that prevailing AD staging schemes are designed primarily to capture the anatomical progression patterns of histopathologically detectable lesions in autopsy. For example, Braak stage and CERAD denotes scores on an ordinal scale that, by definition, index the regional spread of detectable tau tangles and β-amyloid plaques, respectively ^73–75^. In contrast, quantitative measures such as global β-amyloid burden provide continuous estimates summarizing lesion loads across affected brain regions affected ^76^. Importantly, these staging metrics are defined as an aggregate statistic at the donor level, rather than at a single-cell resolution.

Because pathology staging naturally incorporates a spatial component and is assigned per donor, predicting stage for individual cells is not directly meaningful. We therefore reframed the task as estimating the degree to which a cell’s transcriptomic profile is consistent in donors with high- or low-pathology states. To this end, we trained each inference model (logistic classifier) with a binary objective. Given the excellent calibration of logistic regression estimators, we could derive and interpret the model predicted probabilities on a continuous scale of pathology burden, at the single-cell level.

Independent pathology inference models were trained for five markers used in neurology and neurodegeneration (Braak stage, β-amyloid, CERAD, TDP-43, and Lewy body) across nine cell types, yielding in a collection of 45 separate models. The model pipeline included first selecting *H* highly variable genes out of a total of 23,126 genes common between ROSMAP-PFC and SEA-AD (*highly_variable_genes*, flavor = ‘seurat_v3’; scanpy v1.11.2) ^77,78^. The next step involved standardizing counts for each gene across all *N* cells of a given type (*StandardScalar*; scikit-learn v1.7.2). This was followed by PCA transformation keeping *M* components. Finally, a logistic regression model was trained for binary pathology classification.

To select the optimal model configuration (model selection), during training, each model underwent hyperparameter optimization using a rigorous nested cross-validation (CV) (outer fold *k_o_* = 5, inner fold *k_i_* = 5). For all CV splits, stratification was performed at the donor level, such that test sets (outer fold) and validation sets (inner folds) contained cells from donors unseen during model training. In the inner folds, optimal hyperparameter configuration was selected via an exhaustive grid search based on the best ROC-AUC (mean across folds). The grid search had the following considerations: number of highly variable genes (*H* ∈ [2,000, 22,000]), number of principal components (no PCA or *M* ∈ [20, 500] components), L2 regularization strength (*C* ∈ [10^−7^, 10^8^]), class balancing (class_weight ∈ {balanced, None}), pathology thresholds for binarization (Supplementary table 2), and a choice of demographic covariates (sex: male only, female only, combined; dementia status of participants: Alzhiemer’s dementia, other dementia, MCI, or no impairment ^79,80^).

In the outer folds, the optimal hyperparameter configuration from the inner loop was used to evaluate ROC-AUC on a held-out test set of unseen cell observations. The logistic regression model was implemented using *sklearn.linear_model.LogisticRegression* with the ‘saga’ solver and L2 regularization. The maximum number of iterations was set to 500, and all other parameters were left at their default values.

Final hyperparameter combinations were chosen based on their aggregated performance across outer folds, defined as the configuration most frequently selected as optimal in the inner loop. In case of ties, models were selected based on increasing generalizability that prioritized greater separation between predicted classes (higher KL divergence between the predicted probabilities of the high versus low classes). We observed that this usually favored models having a richer feature space (higher highly variable genes), lower latent dimensionality (fewer principal components), and intermediate regularization strengths (Supplementary table 3).

The 45 pathology estimators (5 pathologies, 9 cell types) were retrained 10 times with the optimal hyperparameter configuration and randomly selected subset of cells (50-80%) of the respective cell type in ROSMAP-PFC (*scanpy.pp.subsample*). These models were subsequently applied to external datasets to infer neuropathology scores (mean and std; cf. below).

Note that the model estimated scores (pathology-proxy) represent cell type-resolved transcriptomic similarity to neuropathology markers rather than direct estimates of pathological burden (such as with histology or immunohistochemistry).

### Model level gene contribution

To obtain gene-level effect size for the sake of interpretability, the model weights were translated back to the original ambient gene space via,

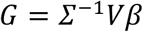

where *β* (*M* components x 1) is the logistic regression derived coefficient vector in the PCA space, V is the loading matrix from PCA (*H* genes x *M* components), and *Σ*^−1^ = diag(*σ*_1_, …, *σ*_H_), such that *σ*_i_ is the gene specific standard deviation (i.e., inverting the standard scaling step). Note that for our analysis, we ignored the estimated intercept parameter and focused on the relative contribution of genes to each model.

To assess the robustness of the predictive contribution of each gene (i.e., its beta coefficient), we refit 500 bootstrapped pathology-estimator models using the optimal hyperparameters for the specific cell type. Specifically, in each bootstrap iteration, ROSMAP-PFC donors (instead of cells) were sampled with replacement to simulate random participant samples from the same population. The full model was refit in each iteration. For each gene, bootstrap percentile intervals were estimated from the resulting weight distribution, and robustness was evaluated based on 5/95 CI, and whether the interval excluded zero. Genes which did not pass the robustness check were excluded from further analysis.

All models and computed metrics were tested for significance using a permutation framework. For each pathology estimator, binary pathology labels were randomly permuted across cells, and the estimator was retrained using the permuted labels (N=1000 iterations). This permutation procedure selectively destroyed the association between the transcription profiles (model input) and the pathology measure (model output), while preserving the inter-gene correlation structure. The resulting null models were then applied to downstream datasets to generate corresponding null distributions of predicted pathology scores (cf. below). These distributions were used to assess the significance of score-derived summary statistics in downstream datasets (Supplementary Fig. 2a, b).

### Cell type inference model

To align cell identities in external datasets with their closest transcriptomic counterparts in the ROSMAP-PFC training dataset, we built a cell type inference model. This alignment step was applied to each cell in downstream datasets (SEA-AD, ROSMAP-MR, *Siletti*) to identify the corresponding ROSMAP-PFC cell type and select the appropriate cell type-specific pathology estimator. Please note that the inferred labels were not used to overwrite existing cell annotations in the external datasets.

We trained a one-vs-rest logistic regression model to distinguish nine major cell types based on whole-transcriptome inputs. Within each cell type, observations were randomly sampled such that 50% of the population was retained. Gene expression values were standardized by z-scoring each gene across all cells (mean-centered and scaled to unit variance). The thus centered rescaled expression matrix was used as input to a supervised classification model. Practically, the model was implemented using *sklearn.linear_model.LogisticRegression* with the ‘saga’ solver and L2 regularization, wrapped in *sklearn.multiclass.OneVsRestClassifier* (scikit-learn v1.7.2). The maximum number of iterations was set to 500, and all other parameters were left at their default values.

For neuronal subtype inference, we trained independent secondary classifiers within each ROSMAP-PFC neuronal cell population. One-vs-rest logistic regression models were trained within excitatory neurons of set 2 (9 subtypes), excitatory neurons of set 3 (6 subtypes), and inhibitory neurons (25 subtypes). Model training followed the same preprocessing and modeling procedure as described for the primary cell type inference model (see above).

### Carrying over pretrained pathology estimation models to downstream datasets

Before application of the pathology estimation pipeline, all downstream datasets (SEA-AD, ROSMAP-MR, *Siletti*) underwent pre-processing identical to that of the ROSMAP-PFC training dataset. For each cell, the total gene transcript count was normalized to 10,000 UMIs. Cells were then assigned to one of the major ROSMAP-PFC cell types using the already trained cell type inference model (cf. above). This provided a data-driven method to determine which ROSMAP-PFC cell type-specific pathology estimation model was most appropriate for each cell.

Single cell-level pathology scores were computed by applying the corresponding pretrained ROSMAP-PFC pathology estimator to each cell’s transcriptome. The original preprocessing and training pipeline was reused without modification. This included selection of the predefined highly variable genes, standardization of gene expressions using the fitted *StandardScalar*, projection into the existing PCA space, and pathology score inference using the trained logistic regression model.

As no single evaluation measure captures all aspects of generalization and its possible failures ^81^, we quantified model performance in SEA-AD cells using multiple complementary metrics — ROC-AUC, AUPRC, and conditional precision (see below). Notably, binarizing SEA-AD pathological marker stages into high- and low-pathology groups resulted in imbalanced cell-level labels, driven by the uneven distribution of donors across pathology stages. Accordingly, we used AUPRC which is more sensitive to class imbalance and better reflects performance on the positive class.

### Subtype specific pathology estimators

To account for biological variability among cell subtypes, we trained pathology estimators at the level of individual neuronal subtypes. The ROSMAP-PFC dataset provided subtype information for excitatory neurons of set 3 (nine subtypes), and excitatory neurons of set 2 (four subtypes), and were considered for this extended method. Since our comprehensive validations in the SEA-AD dataset were with the Braak and Abeta models, pathology models for only these two markers were trained and evaluated with the subtype trained models.

To estimate subtype specific pathology, we followed the same pipeline as before. First, the cell subtype inference model trained on ROSMAP-PFC (see above) was applied to assign subtype labels to cells previously inferred as excitatory neurons of set 3 in the downstream datasets. Next, subtype specific pathology estimators (trained independently in ROSMAP-PFC for each subtype) were used to estimate single-cell level pathology.

During training, the previously described nested CV procedure was applied with the same hyperparameter search space. Training and evaluation were restricted to the male cohort (cf. above).

### Conditional precision

To obtain a calibration agnostic, comparable summary across datasets, we computed conditional precision for each pathology estimator. Although ROC-AUC and AUPRC summarize ranking performance across score thresholds, they are difficult to compare across datasets when score distributions, class prevalence, or calibration differ. Here, we therefore asked whether the highest scoring cells from each estimator were enriched for cells from the diseased donors. Conditional precision was defined as the fraction of correctly classified cells among those with the highest model-assigned pathology probabilities. Specifically, in downstream validation datasets (SEA-AD and ROSMAP-MR), we computed the 50^th^ to 99^th^ percentiles of the predicted pathology scores at fixed percentile intervals for each ROSMAP-PFC cell type. Formally, let *P* denote the set of all cells of a given cell type. Then, *P*(t) ⊂ *P* such that cells in *P*(t) have model probabilities greater than or equal to the *t*^th^ percentile of scores. The conditional precision for *P*(*t*) was computed as,

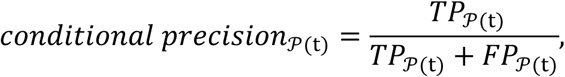

where *TP_P_*_(t)_ and *FP_P_*_(t)_ denote the number of true and false positives among cells in *P*(t), respectively. Because this metric evaluates precision using score rankings rather than raw probability values, it remains robust to shifts in score distributions.

For comparison, the baseline precision was computed as *Precision_P_* = *TP_P_*/*N*, where *TP_P_* is the number of true positives and *N* is the total number of cells in *P*. This corresponds to the expected precision under random selection.

### SEA-AD and ROSMAP multi-region dataset

To quantify the pathology prediction performance of the ROSMAP-PFC trained models, we used the SEA-AD and ROSMAP-MR snRNA-seq datasets. As ground truth, we used the gold standard reference pathology annotations provided in these datasets.

Within the neocortical setting of SEA-AD, binarization of reference pathology was performed using explicit thresholds on donor-level markers. These thresholds were defined from the best models derived in ROSMAP-PFC (see hyperparameter search above). This high vs low classification enabled subsequent evaluations with ROC-AUC and AUPRC.

For evaluations of the models in ROSMAP-MR, we considered the inherent regional component of the pathology staging definitions. For Braak stage, defined by spatially ordered progression of detectable neurofibrillary tangles, there is no biologically justified way to impose a single donor-level binarization that applies consistently across different brain regions. Similarly, β-amyloid is a brain-wide summary of amyloid plaque burden, and a single binarization score is arbitrary. Therefore, for cross-region performance evaluations, we used rank-based metrics that directly related model scores to region-specific reference pathology.

For assessment of ROSMAP-MR donor-level estimates, an aggregated score was computed for each donor, per region and pathology. This metric was defined as the 90^th^ percentile value from the score distributions of cells of a given inferred type, averaged across all evaluated cell types ^(^*^q^*0.9,avg^).^

Associations between model-derived pathology-estimates and reference pathology was computed using Kendall’s tau-b correlation metric. Specifically, the metric was evaluated between region-and cell type-specific estimated pathology scores and donor-region-level reference pathology. We used the AG, EC, HC, MT and PFC specific reference donor pathologies for neurofibrillary tangles and β-amyloid.

### Downstream gene contributions

Previously, we derived gene effect sizes for pathology prediction at the level of ROSMAP-PFC cell types. However, in downstream datasets, the native cell population annotations did not always correspond uniquely to a ROSMAP-PFC cell type using our cell type inference model. More importantly, the baseline gene expression signatures of novel cell populations in downstream datasets differ substantially from their closest ROSMAP-PFC counterparts.

This difference in baseline gene expression can also be present within a major cell population that is sampled from different brain regions. Indeed, prior cross-region analyses have shown that the cell types have substantial diversity across the brain. This diversity is conserved in both humans and mice, giving rise to transcriptionally distinct regional cell populations ^17,82^. Consequently, direct interpretation of gene coefficients is confounded by brain regions, even within the same major cell populations.

To account for these differences and homogenize the attribution of genes to specific supercluster and brain regions, we employed a two-step approach. In the first step, we used a Shapley-based attribution framework ^83^, decomposing individual gene contributions to pathology prediction at the resolution of single cells (see below). These per-cell gene attributions could then be summarized using aggregate statistics at any arbitrary resolutions, such as brain region and supercluster.

In the second step, we compared the SHAP-derived gene contributions between high- and low-scoring cells within each population of interest. These comparisons were performed using a statistical framework designed to capture systematic shifts in contribution patterns (see below). This procedure corrected for residual baseline effects and accommodated potential inversion of gene contributions across regions or cell populations, thereby prioritizing genes whose contributions were enriched in high-scoring cells.

#### Step 1: Single-cell gene contribution - SHAP

Exact Shapley contributions are intractable for our setting, as the model operates on ∼2000 input genes. However, for linear models, Shapley values have a closed-form analytical solution. This linearity must hold in both input and output spaces. Our classifier is linear in the logit output space but the PCA transformation mixes gene-level signals in the input space. Consequently, gene-level attributions in the original feature space are no longer independent, rendering direct application of the closed-form Shapley solution inappropriate for our model.

We therefore adopted an interventional SHAP formulation to compute Shapley values in a computationally tractable manner. Under this framework, we quantified the effect of perturbing individual genes on model predictions in the logit space, while treating other genes as independently varying. This interventional approach is analogous to differential gene expression analyses, which assess gene-wise associations with a phenotype despite underlying correlation structure. Under this independence formulation, the SHAP value for a gene *j* in cell *i* from a specific population of interest *P* was computed as,

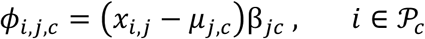

where *x_i, j_* denotes the expression of gene *j* in cell *i*; *β_j_* is the back-projected model coefficient for gene *j* from ROSMAP-PFC cell type *c*; *P*_c_ denotes the set of cells inferred as *c* and belonging to *P*; *μ_j,c_* is the median expression of gene *j* in *P*_c_. Note that the contributions were median-centered, instead of the SHAP derived analytical baseline of mean-centering. This consideration is consistent with common practices in single-cell analysis accounting for skewed distributions of gene expressions ^78,84^.

#### Step 2: Aggregating gene contribution within a population of interest – differential SHAP

For each gene, we compared SHAP-derived contribution distributions between high- and low-pathology-proxy cells. The magnitude of the difference was quantified using Wasserstein distance, a parameter-free metric sensitive to global shifts in distribution rather than changes in summary statistics alone. The direction of the shift was assessed using Cohen’s *d*, and statistical significance was evaluated using a two-sided Mann-Whitney U test. Resulting *p*-values were corrected for multiple testing using the Benjamini-Hochberg FDR procedure, applied independently across all genes within each supercluster and each of the ten major brain regions.

The Wasserstein distance, 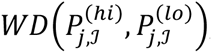, was computed for each gene *j* and population of interest *P*. We defined the high and low empirical SHAP distributions 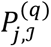 as,

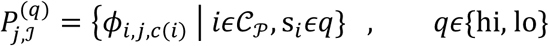

where, *c(i)* denotes the inferred cell type of cell *i*; 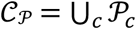 denotes the set of all cells belonging to *P*, across all inferred cell types; *s_i_* is the model score for cell *i*. Here, *q* = hi denotes cells with pathology proxy scores 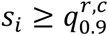, and *q* = lo denotes cells with 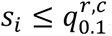, where quantiles are computed for each brain region *r* and ROSMAP-PFC cell type *c*.

### Filtering datasets

#### General filtering

For all downstream datasets, first, all cells inferred as vascular cells were excluded given poor model performance of this class across all pathology markers (see Results). Next, cells with the second highest class probability greater than a heuristic threshold were discarded as having a confused classification by the model. The threshold was determined after a combined examination of the probability distributions of the second highest class probabilities and minimizing the total number of cells that would be excluded given a threshold.

#### Siletti specific filtering

The cells in *Siletti* were examined to identify potential outlier cells. These cells were removed to ensure that downstream results represented robust biological patterns rather than sampling artifacts or misclassified samples. We identified outliers as follows,

i. We filtered out nuclei using the general filtering described, resulting in exclusion of all choroid plexus and ependymal cells in this step.
ii. Inferred cell types contributing less than 10% of all nuclei within a *Siletti* defined supercluster×region of interest combination was filtered out. This step removed spurious cell type inferences. For example, 89 out of 66,837 hippocampal dentate gyrus nuclei inferred as astrocytes were removed (Supplementary table 17).
iii. We removed superclusters from certain brain regions where their relative abundance fell below the 30^th^ percentile across all regions or contained fewer than 100 nuclei in the supercluster×region combination. For example, 687 medium spiny neurons from the cerebral cortex were filtered out.

Together, these filtering steps enabled retention of only well-represented populations in the *Siletti* dataset. In total, 152,189 cells were filtered out across all brain regions, retaining 2.89 million cells for downstream analysis.

#### Relative regional contribution

Regional contributions to predicted pathology burden were computed by summing predicted pathology scores across all cells within each region, aggregating across cell types and donors. For each region, this summed score was then divided by the total summed pathology score across all cells from the dataset.

#### Pseudo-vulnerable populations

The commonly used approach for identifying vulnerable cell populations compares the differential abundance of a given population between control and disease groups. However, neuronal vulnerability in AD is not solely reflected by cell loss. Neuropathological studies have shown that neurons containing aggregated tau can persist for decades while exhibiting substantial functional impairment ^60^. In our setting, we therefore do not rely on case-control differences in cell abundance. Instead, we estimate a pathology proxy at the single cell level based on transcriptomic signatures. To capture this phenomenon, we define a molecular analogue of vulnerability which quantifies the propensity of a cell population to exhibit pathology-like transcriptomic states.

Here, we defined molecular vulnerability of a population of interest (example cell subcluster or brain region) as the difference between representative pathology estimate and that of its corresponding parent population. Formally, for a child population of interest *P* (example cell subcluster) belonging to a parent population *G* (supercluster), molecular vulnerability was computed as,

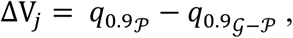

where 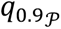 denotes the 90^th^ percentile of score distribution in population *P* and 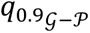 denotes the 90^th^ percentile of the score distribution in *G* after excluding *P*. Significance of the differences were assessed using the appropriate statistical tests and were corrected for multiple testing across all features within the corresponding comparison group (e.g., all subpopulations or all brain regions) using the Benjamini-Hochberg correction procedure ^85^. Only populations of interest with more than 50 observations were considered for this analysis.

“Pseudo-vulnerable” populations were populations with statistically significant molecular vulnerability. We used this approach to identify pseudo-vulnerable cell subpopulations in SEA-AD, and pseudo-vulnerable cell subpopulations and brain regions in *Siletti*.

### Region ranking

To rank the 108 brain regions in *Siletti* by inferred pathology burden we computed each region’s molecular vulnerability as defined above.

Specifically, model-derived pathology scores derived at the single-cell level were first normalized within each supercluster. Normalization was performed by subtracting the median score computed across all donors and regions. The normalized scores were then used to compute molecular vulnerability of brain regions. Specifically, all cells from a region were defined as the population of interest *P* and remaining cells formed *G*. Statistical significance was assessed using a two-sided Mann-Whitney U test, and the *p*-values were corrected for multiple testing across regions using the Benjamini-Hochberg procedure.

For per-donor and inferred cell type-specific analyses, the same procedure was applied separately within each donor or inferred cell type. Comparison between ranked regions was performed using a weighted measure of Kendall’s tau ^86^, which assigns greater weight to concordance among the highest-scoring regions (implemented using *scipy.stats.weightedtau;* SciPy v1.15.2).

### Alternate region ranking

To validate the robustness our primary rankings (see above), we employed an alternative analytical approach to rank the 108 brain regions. We fitted an additive linear model with predicted single-cell pathology scores as the response variable and supercluster and major brain-region as categorical covariates. Residuals from this model were interpreted as pathology scores adjusted for supercluster composition and were subsequently used for the region ranking. We note that this approach is more restrictive, as it removes regional variation in supercluster composition and may therefore attenuate biologically meaningful differences.

### Brain Map

To visualize dissection-level pathology estimates, we mapped the 2D *Siletti* dissections (following the Allen Human coronal reference atlas ^87^) to 84 regions in the 3D Allen Human Reference Atlas. The mapping was taken from previously curated expert-defined mappings ^23^. Atlas NIfTI volumes were downloaded from the Allen institute for brain science [resource](https://download.alleninstitute.org/informatics-archive/allen_human_reference_atlas_3d_2020/version_1/examples/flip_annotation/). The region-wise one-versus-rest pathology scores computed previously (FDR<0.05; see above) were assigned to their corresponding atlas parcels. Each atlas parcel was uniformly filled with its associated score to generate 3D volumetric statistical maps in atlas space. The statistical maps were visualized using Nilearn v0.12.1 (*plot_stat_map*).

### Supercluster contribution

To quantify the relative contribution of each *Siletti* cluster (423 cell sub-populations) to inferred pathology scores, within each brain region we first identified high pathology cells. These cells were defined as those exceeding the 90^th^ percentile of model-derived pathology scores within that region. For each cluster, we computed the fraction of its cells present among these high-pathology cells, which served as a quantitative measure of cluster contribution. Clusters with fewer than 20 cells within a given brain region were excluded from analyses for that region.

To account for the difference in the number of cell observations from each of the 423 clusters, statistical significance of enrichment was assessed using a one-sided hypergeometric test. In other words, this test quantified the significance of observing a given number of high-scoring cells within each cluster, given the total number of cells in that cluster, the overall number of high-scoring cells across all clusters, and the total number of cells across all clusters.

Concretely, we computed *p*-values of enrichment using the survival function of the hypergeometric distribution, testing for over-representation as,

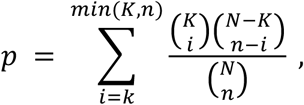

where, *N* is the total number of cells in the region, *K* is the number of cells in a cluster from that region, *n* is the number of high pathology cells, and *k* is the number of cluster cells among the high pathology cells. Multiple testing correction was performed across all clusters and brain regions using the Benjamini-Hochberg procedure.

### Layer inference model

To infer cortical layer identity (i.e., L1 – L6 and white matter) from transcriptomic profiles, in datasets that do not provide this kind of annotation for neocortical cells, we designed a hierarchical two-step classification framework. As a preparatory steps, cells were first classified as white matter versus grey matter. Cells predicted as grey matter were subsequently assigned to one of six neocortical layers.

Two classifiers were trained for these tasks. The white-versus-gray matter binary classifier was trained on annotated snRNA-seq data from *Maynard* ^33^. The six-layer classifier (model below) was trained on annotated data from *Jorstad* ^32^. Both models used identical pre-processing pipelines. Raw gene expression counts were normalized at the cell level as described above. The top 5,000 highly variable genes were selected from the dataset. Analysis was restricted to only protein coding and long-noncoding RNA genes. Gene expression counts were then z-scored across all cells (scikit-learn v1.7.2; *StandardScalar*).

The inference models were trained using sklearn *LogisticRegression* in a one-versus-rest classification scheme (scikit-learn v1.7.2, solver = ‘saga’; regularization strength C = 0.001; class_weight = ‘balanced’; remaining parameters default). Qualitative model evaluation was based on concordance between predicted layer assignments and established biological expectation in downstream datasets (Figure 7b). For the white matter-grey matter classifier, model performance was evaluated by its application on the *Jorstad* dataset. The average accuracy was 0.79±0.34 across cell subclasses (Supplementary Fig. 8c). For the six-layer classifier, quantitative model performance was evaluated using leave-one-donor-out CV to prevent donor-specific information leakage. Within each fold, models were trained on all but one donor and evaluated on the held-out donor (Supplementary Fig. 8d).

### Regulatory network analysis

Gene regulatory networks (GRNs) were inferred separately for each *Siletti* supercluster using pySCENIC (v0.12.1) ^88^ with default parameters. As described in the pySCENIC workflow, transcription factor (TF)–target co-expression modules were first identified using the *GRNBoost2* algorithm ^89^. The full list of human transcription factors was obtained from the Aerts lab resource (https://resources.aertslab.org/cistarget/tf_lists/). Subsequently, GRN modules were pruned by cis-regulatory motif enrichment analysis using the human motif database (v9; https://resources.aertslab.org/cistarget/motif2tf/). This retained biologically plausible TF-target interactions supported by motif evidence.

For each TF, the regulon was defined as the TF and its predicted target genes retained after motif enrichment filtering. When multiple motif annotations were present, the regulon with the highest normalized enrichment score was selected. Regulon activity was then quantified at the single-cell cell resolution using *AUCell* (pySCENIC v0.12.1), yielding continuous activity score. The resulting regulon set encompassed the full spectrum of cellular programs represented in the dataset, including baseline homeostatic functions.

To relate regulon activity to the spatial distribution of estimated pathology, associations between per-cell regulon activity and pathology load were quantified using mutual information (MI). MI was estimated with *mutual_info_regression* function from scikit-learn (v1.7.2; default parameters). Because MI is non-directional, the direction of association was determined using Kendall’s tau-b correlation. Thus, the regulons with positive MI were preferentially activated in high-pathology contexts, whereas those with negative MI were suppressed.

Statistical significance was assessed by permuting pathology labels across cells while keeping regulon activity fixed (*n*=1000 iterations). Regulons with observed signed MI values exceeding the 95^th^ percentile or below the 5^th^ percentile of the permutation distribution were considered significant.

### CSF protein markers

We assessed concordance between SHAP-derived gene contributions and inter-individual differences in CSF proteomic markers associated with early AD. CSF protein markers were obtained from a previously published cohort of 1,760 participants spanning the AD spectrum ^27^. Preclinical AD was defined as cognitively normal individuals with abnormally decreased CSF β-amyloid levels. The study reported proteins differentially detected between preclinical AD cases with elevated total tau and control individuals. In total, 2,535 CSF proteins were measured. Of these, 1,446 proteins overlapped with genes profiled in our transcriptomic analysis (1,446/23,126). This intersecting set of genes was used as the background for enrichment testing. Overlap enrichment was evaluated using a one-sided hypergeometric test.

### Gene ontology and pathway enrichment analysis

Functional enrichment analysis for pathology associated regulons was performed using the gseapy *enrichr* package (v1.1.9; accessed November 2025). Specifically, for each supercluster-specific regulon (see above), gene sets were tested for over-representation against curated gene set databases available through the Enrichr platform (https://maayanlab.cloud/Enrichr/#libraries). We examined GO_Biological_Processes_2023, GO_Molecular_Function_2023, KEGG_2021_Human, and SynGO_2024 terms. Enrichment significance was assessed using the *Enrichr* over-representation framework which provides p-values corrected for multiple-testing. For illustration purposes, we curated biological topics relevant to AD based on prior literature ^5,21^; enriched terms were subsequently mapped to these curated topics.

## Acknowledgements

ROSMAP is supported by P30AG10161, P30AG72975, R01AG15819, R01AG17917, U01AG46152, and U01AG61356. ROSMAP resources can be requested at https://www.radc.rush.edu. DB was supported by the Brain Canada Foundation, through the Canada Brain Research Fund, with the financial support of Health Canada, National Institutes of Health (NIH R01 AG068563A, NIH R01 DA053301-01A1, NIH R01 MH129858-01A1), the Canadian Institute of Health Research (CIHR 438531, CIHR 470425), the Healthy Brains Healthy Lives initiative (Canada First Research Excellence fund), the IVADO R3AI initiative (Canada First Research Excellence fund), and by the CIFAR Artificial Intelligence Chairs program (Canada Institute for Advanced Research).

## Ethics declarations

### Data and Code Availability

All code will be made publicly available on GitHub upon publication of the manuscript and is available from the corresponding author upon reasonable request.

## Extended Data

**Table 1.** Abbreviations.

| Category | Abbreviation | Full Term |
| --- | --- | --- |
| Siletti superclusters | AmygEx | Amygdala excitatory |
| Siletti superclusters | Ast | Astrocyte |
| Siletti superclusters | Ber | Bergmann glia |
| Siletti superclusters | CGEInter | CGE interneuron |
| Siletti superclusters | COpc | Committed oligodendrocyte precursor |
| Siletti superclusters | CerebIn | Cerebellar inhibitory |
| Siletti superclusters | ChoPle | Choroid plexus |
| Siletti superclusters | DLCn6b | Deep-layer corticothalamic and 6b |
| Siletti superclusters | DLI | Deep-layer intratelencephalic |
| Siletti superclusters | DLNP | Deep-layer near-projecting |
| Siletti superclusters | EMSN | Eccentric medium spiny neuron |
| Siletti superclusters | Epen | Ependymal |
| Siletti superclusters | Fib | Fibroblast |
| Siletti superclusters | HCA13 | Hippocampal CA1-3 |
| Siletti superclusters | HiCA4 | Hippocampal CA4 |
| Siletti superclusters | HiDG | Hippocampal dentate gyrus |
| Siletti superclusters | LLCh | LAMP5-LHX6 and Chandelier |
| Siletti superclusters | LRL | Lower rhombic lip |
| Siletti superclusters | MB | Mammillary body |
| Siletti superclusters | MDIn | Midbrain-derived inhibitory |
| Siletti superclusters | MGEInter | MGE interneuron |
| Siletti superclusters | MSN | Medium spiny neuron |
| Siletti superclusters | Mic | Microglia |
| Siletti superclusters | Misc | Miscellaneous |
| Siletti superclusters | Oli | Oligodendrocyte |
| Siletti superclusters | Opc | Oligodendrocyte precursor |
| Siletti superclusters | Spl | Splatter |
| Siletti superclusters | ThEx | Thalamic excitatory |
| Siletti superclusters | ULInter | Upper-layer intratelencephalic |
| Siletti superclusters | URL | Upper rhombic lip |
| **Siletti superclusters** | Vasc | Vascular |
| **Major regions** | CC | Cerebral cortex |
| **Major regions** | CN | Cerebral nuclei |
| **Major regions** | EP | Epithalamus |
| **Major regions** | HB | Hindbrain |
| **Major regions** | HC | Hippocampus |
| **Major regions** | HT | Hypothalamus |
| **Major regions** | MB | Midbrain |
| **Major regions** | PC | Paleocortex |
| **Major regions** | SC | Spinal cord |
| **Major regions** | TH | Thalamus |
| **RADC-variable** | braak_stage | Braak stage |
| **RADC-variable** | amyloid | Amyloid |
| **RADC-variable** | tdp_st4 | TDP-43 |
| **RADC-variable** | dcfdx | Clinical cognitive diagnosis summary |
| **RADC-variable** | dlbdx | Lewy body |
| **RADC-variable** | ceradsc | CERAD score |
| **RADC-variable** | age_death | Age at death |
| **Others** | CERAD | Consortium to Establish a Registry for Alzheimer's Disease |
| **Others** | MCI | Mild cognitive impairment |
| **Others** | FDR | False discovery rate |
| **Others** | OR | Odds ratio |
| **Others** | GABA | Gamma-aminobutyric acid |
| **Others** | MAPK | Mitogen-activated protein kinase |
| **Others** | SEA-AD | Seattle Alzheimer's Disease Brain Cell Atlas |
| **Others** | ASAP | Aligning Science Across Parkinson's |
| **Others** | SynGO | Synaptic Gene Ontologies |
| **Others** | TDP-43 | Transactive response DNA-binding protein 43 kDa |
| **Others** | VGLUT1/2/3 | Vesicular glutamate transporter types 1, 2, and 3 |

